# Molecular determinants of β-arrestin coupling to formoterol-bound β_1_-adrenoceptor

**DOI:** 10.1101/2020.03.27.011585

**Authors:** Yang Lee, Tony Warne, Rony Nehmé, Shubhi Pandey, Hemlata Dwivedi-Agnihotri, Patricia C. Edwards, Javier García-Nafría, Andrew G.W. Leslie, Arun K. Shukla, Christopher G. Tate

## Abstract

The β_1_-adrenoceptor (β_1_AR) is a G protein-coupled receptor (GPCR) activated by the hormone noradrenaline, resulting in the coupling of the heterotrimeric G protein G_s_^1^. G protein-mediated signalling is terminated by phosphorylation of the receptor C-terminus and coupling of β-arrestin 1 (βarr1, also known as arrestin-2), which displaces G_s_ and induces signalling through the MAP kinase pathway^2^. The ability of synthetic agonists to induce signalling preferentially through either G proteins or arrestins (biased agonism)^3^ is important in drug development, as the therapeutic effect may arise from only one signalling cascade, whilst the other pathway may mediate undesirable side effects^4^. To understand the molecular basis for arrestin coupling, we determined the electron cryo-microscopy (cryo-EM) structure of the β_1_AR-βarr1 complex in lipid nanodiscs bound to the biased agonist formoterol^5^, and the crystal structure of formoterol-bound β_1_AR coupled to the G protein mimetic nanobody Nb80^6^. βarr1 couples to β_1_AR in a distinct manner to how G_s_ couples to β_2_AR^7^, with the finger loop of βarr1 occupying a narrower cleft on the intracellular surface closer to transmembrane helix H7 than the C-terminal α5 helix of G_s_. The conformation of the finger loop in βarr1 is different from that adopted by the finger loop in visual arrestin when it couples to rhodopsin^8^, and its β-turn configuration is reminiscent of the loop in Nb80 that inserts at the same position. β_1_AR coupled to βarr1 showed significant differences in structure compared to β_1_AR coupled to Nb80, including an inward movement of extracellular loop 3 (ECL3) and the cytoplasmic ends of H5 and H6. In the orthosteric binding site there was also weakening of interactions between formoterol and the residues Ser211^5.42^ and Ser215^5.46^, and a reduction in affinity of formoterol for the β_1_AR-βarr1 complex compared to β_1_AR coupled to mini-G_s_. These differences provide a foundation for the development of small molecules that could bias signalling in the β-adrenoceptors.

---

Ligand bias arises through differential activation of the G protein pathway compared to the arrestin pathway and has been observed in ligands binding to many different GPCRs such as the μ-opioid receptor^9^, the angiotensin receptor (AT_1_R)^10^ and the β-adrenoceptors, β_1_AR and β_2_AR^5,11^. The ligands can show complex pharmacology. For example, carvedilol is an inverse agonist of β_1_AR when G protein activity is measured, but it is a weak agonist of the MAPK pathway activated by βarr1^11^. In contrast formoterol is an agonist of both pathways, but stimulates the βarr1 pathway more than the G protein pathway^5^. Current theories favour the hypothesis that GPCRs exist in an ensemble of conformations and that ligands preferentially stabilise specific conformations^12^. This suggests that the conformation of a receptor bound to a heterotrimeric G protein could be different from the conformation that binds βarr. There is considerable spectroscopic evidence to support the existence of an ensemble of conformations of a GPCR even in the absence of ligands, and that specific ligands selectively stabilise specific conformations^13-15^. A notable study on AT_1_R using electron paramagnetic resonance also supports the notion that βarr biased ligands stabilise a different subset of conformations compared to G protein biased ligands^16^. Structures have been determined of many different GPCRs and a number of distinct states have been identified. For example, X-ray crystallography has identified two major structural states of β-adrenoceptors, comprising a number of inactive states^17-19^ and a number of active states^6,7,20-22^. In the absence of a G protein, the most thermodynamically stable state of the agonist-bound receptor is very similar to the inactive state, except for a small contraction of the ligand binding pocket. The active states have to be stabilised by binding of a G protein or a G protein-mimetic (*e.g.* nanobody Nb80 or Nb6B9)^6,22^ at the intracellular cleft that opens transiently on the cytoplasmic face of the receptor with increased frequency upon agonist binding^13^. Although the details of how an agonist activates the β-adrenoceptors is known in great detail^6,7,17^ and also the molecular basis for how G protein coupling increases ligand affinity^21^, there are few molecular details alluding to the mechanism of biased agonism. The structure of carvedilol-bound β_1_AR has been determined in the inactive state, and this suggested that interactions of the ligand with the extracellular end of H7 could promote the biased effect of the ligand as this is not observed in other unbiased ligands^23^. However, an active-state structure of β_1_AR coupled to βarr1 is required to further understanding of biased signalling, so we determined the cryo-EM structure of the formoterol-bound β_1_AR-βarr1 complex.

A complex of βarr1 coupled to purified β_1_AR could not be formed effectively in detergent, so β_1_AR was inserted into nanodiscs (see Methods). The β_1_AR construct contained six mutations to improve thermostability and a sortase sequence to allow ligation of a phosphorylated peptide (V_2_R_6P_) identical to the C-terminal sequence of the vasopressin receptor V_2_R. Pharmacological analysis of purified β_1_AR in nanodiscs indicated that coupling of βarr1 caused an increase in agonist affinity (Extended Data Fig. 1) as observed for coupling of mini-G proteins to detergent-solubilised β_1_AR^21^. This implied that the receptor coupled to arrestin was in an active state, as has been also observed for other GPCRs^24^. The structure of the formoterol-bound β_1_AR-βarr1 complex in nanodiscs (Fig. 1a-c) was determined by cryo-EM (see Methods and Extended Data Figs. 2-5), in the presence of the antibody fragment F_ab_30 that locks arrestin into an active conformation^25^ and was essential to provide sufficient mass for the alignment of the particles. The overall resolution was 3.3 Å, with the best-resolved regions of the cryo-EM map at the interface between β_1_AR and βarr1, reaching a local resolution of 3.2 Å (Extended Data Fig. 2 and Table 1). We also determined the structure of formoterol-bound β_1_AR-Nb80 complex in detergent by X-ray crystallography at 2.9 Å resolution (Fig. 1d and Table 2) to allow a direct comparison between β_1_AR coupled to different proteins but bound to the same ligand. The β_1_AR-Nb80 complex is too small (∼50 kDa of ordered protein) for structure determination by single-particle cryo-EM. The overall structures of formoterol-bound β_1_AR in the β_1_AR-Nb80 and β_1_AR-βarr1 complexes were virtually identical (rmsd 0.6 Å).

**Fig. 1.**
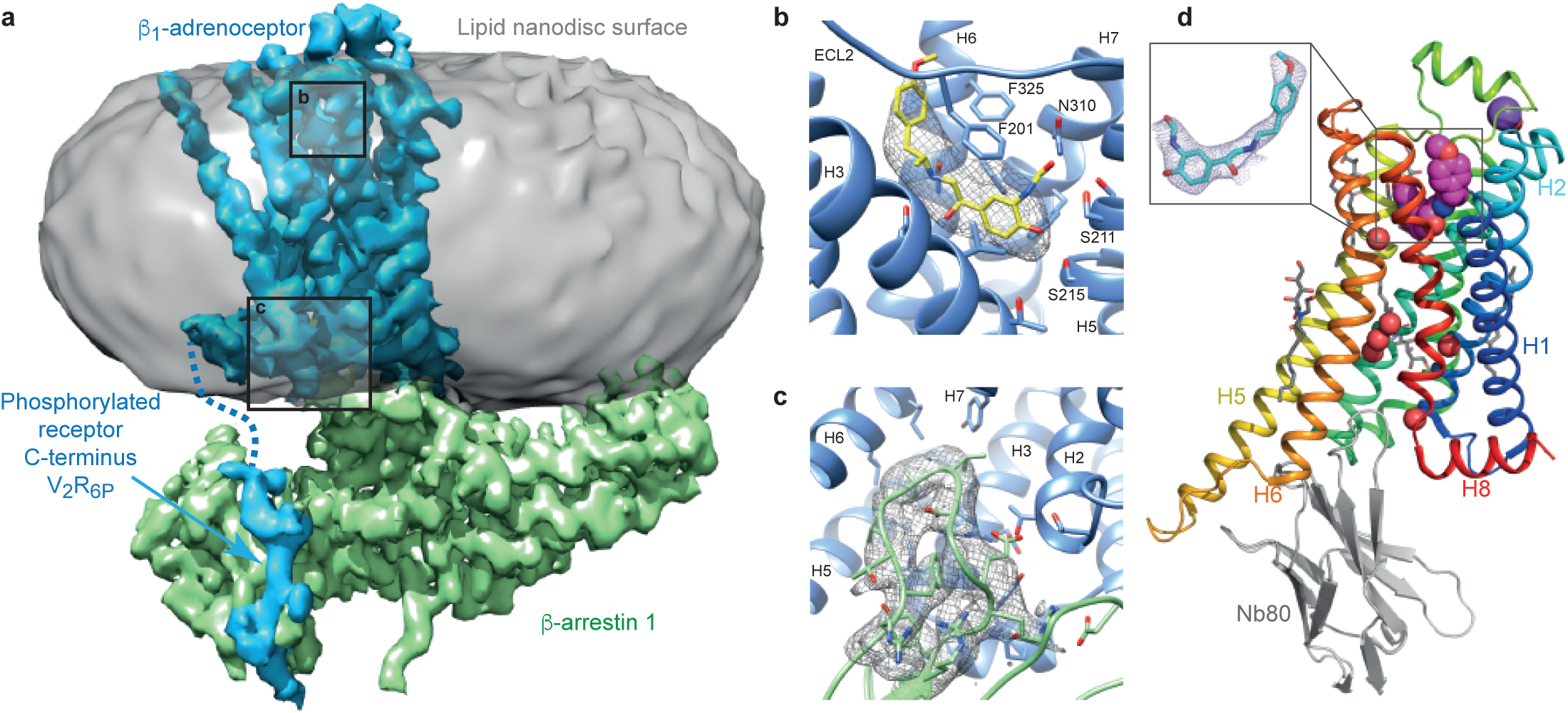
Overall cryo-EM reconstruction of the β_1_AR-βarr1 complex. **a**, The density of the cryo-EM map (sharpened with a *B* factor of −80 e/Å^2^) is coloured according to polypeptides (β_1_AR, blue; βarr1, green) and overlaid on density of the nanodisc (grey). F_ab_30 has been omitted from the structure for clarity (see Extended Data Fig 2). **b,** The orthosteric binding pocket of β_1_AR (pale blue) with formoterol (sticks: yellow, carbon) and its density in the cryo-EM map (grey mesh). **c,** The finger loop of βarr1 with side chains shown as sticks (light green, carbon) and its density in the cryo-EM map (grey mesh). Helix 8 of β_1_AR has been removed for clarity. Maps were contoured at 0.02 (2 Å carve radius in panels **b** and **c**) and visualised in Chimera. **d**, Crystal structure of the β_1_AR-Nb80 complex: β_1_AR, rainbow colouration; Nb80, grey; formoterol, magenta spheres (carbon); water molecules, red spheres; purple sphere, Na^+^ ion; detergent Hega-10, grey sticks (carbon). The inset shows an omit map of formoterol in the β_1_AR-Nb80 complex contoured at 1 σ (blue mesh).

**Fig. 2.**
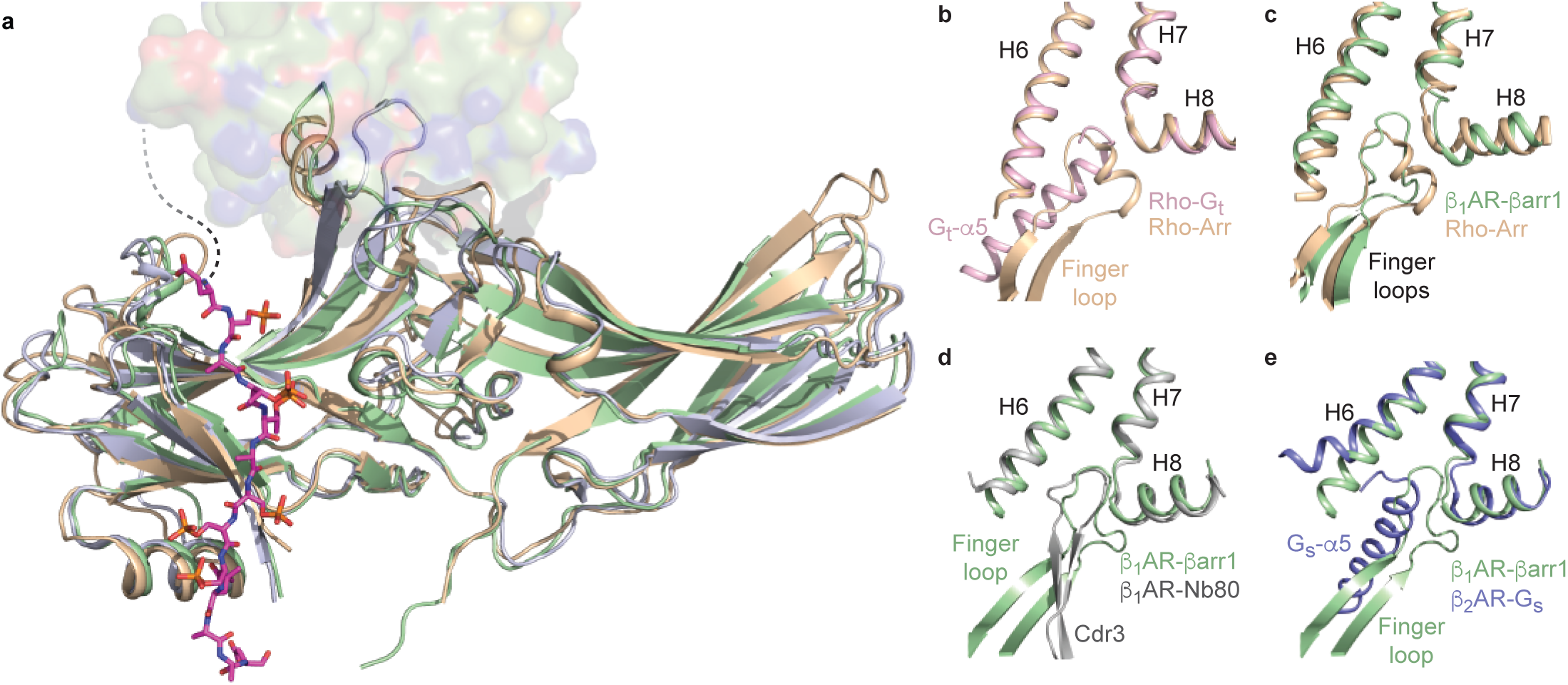
Structure of βarr1 in complex with β_1_AR. **a**, βarr1 (pale green) coupled to β_1_AR (surface representation) was aligned with the structures of S-arrestin (pale brown) coupled to rhodopsin (PDB 5W0P) and the structure of active βarr1 (mauve) bound to the phosphopeptide V_2_Rpp and F_ab_30 (PDB 4JQI). The phosphopeptide shown (carbon, magenta) is V_2_R_6P_ linked to β_1_AR. Full alignments of the phosphopeptides are shown in Extended Data Fig. 6b. **b-e**, details of coupled arrestin finger loops and G protein α5 helices after alignment of the following receptors (PDB code in parentheses) using GESAMT (ccp4 program suite): S-arrestin coupled to rhodopsin (5W0P, pale brown); transducin (G_t_) coupled to rhodopsin (6OYA, pale pink); βarr1 coupled to β_1_AR (pale green); Nb80 coupled to β_1_AR (6IBL, grey); G_s_ coupled to β_2_AR (3SN6, blue).

However, there were small significant differences in the extracellular surface, intracellular surface and in the orthosteric binding site (discussed below). β_1_AR in complex with βarr1 was also very similar to other structures of β_1_AR in an active state (rmsds ∼0.6 Å) and also to β_2_AR coupled to heterotrimeric G_s_ (PDB 3SN6; rmsd 0.8 Å)^7,21^. This allowed a detailed comparison between β_1_AR-βarr1, β_2_AR-G_s_ and β_1_AR-Nb80 (see below).

The overall structure of βarr1 coupled to β_1_AR (Fig. 2) is very similar to the X-ray structure of S-arrestin (also known as visual arrestin) coupled to rhodopsin^8^ (PDB 5W0P; rmsd 1.3 Å, 1853 atoms) and the structure of activated βarr1 coupled to F_ab_30 and the V_2_Rpp peptide^25^ (PDB 4JQI; rmsd 1.1 Å, 1861 atoms). The buried surface area of β_1_AR that makes contact to βarr1 (∼1200 Å^2^; excluding the phosphopeptide interface) is slightly smaller than the surface area of rhodopsin making contact to S-arrestin (∼1400 Å^2^). In addition, there is a 20° difference in tilt of arrestin relative to rhodopsin compared to β_1_AR (Extended Data Fig. 6). However, the regions of β_1_AR and rhodopsin that make contact to either βarr1 or S-arrestin respectively are conserved, as are the positions on the arrestin molecules that make contacts to the receptors (Extended Data Fig. 7 and 8). The position of the C-terminal V_2_R_6P_ peptide in the cryo-EM structure is also virtually identical to the position of the peptide in the crystal structure of the βarr1-F_ab_30-V_2_Rpp complex (Extended Data Fig. 6), with the exception of the phosphate on Thr359. Phospho-Thr359 does not make contacts to βarr1 in the βarr1-F_ab_30-V_2_Rpp complex, but it appears to make contact with the tip of the lariat loop (Lys294, His295) of βarr1 in the β_1_AR-βarr1 complex. No density was observed in the β_1_AR-βarr1 cryo-EM structure equivalent to the N-terminal region of V_2_Rpp (RTpPPSpLGP) that is adjacent to the finger loop in the βarr1-F_ab_30-V_2_Rpp structure; this would clash with the new orientation of the finger loop and the receptor in the β_1_AR-βarr1 complex. The most significant difference between these three structures is the orientation and structure of the finger loop region (Fig. 2). In the activated non-receptor-bound βarr1-F_ab_30-V_2_Rpp structure, the finger loop forms an unstructured region that does not superpose with either the finger loop in S-arrestin coupled to rhodopsin or with βarr1 coupled to β_1_AR. In contrast, the receptor-bound finger loop of S-arrestin and βarr1 superpose, but they adopt different structures (Fig. 2). The finger loop of S-arrestin contains a short α-helical region whereas in βarr1 it forms a β-hairpin. When the arrestin molecules are aligned, it also appears that the tip of the β-hairpin of βarr1 protrudes about 5 Å deeper into the receptor than the α-helical region of S-arrestin. An interesting observation is that the CDR3 loop of nanobody Nb80 that inserts into the receptor bears a remarkable resemblance to the finger loop of βarr1 (Fig. 2), although the polypeptides run in antiparallel directions and bear little sequence similarity except for the Val-Leu residues at the tip of the loops.

Structure determination of the β_1_AR-βarr1 complex in nanodiscs showed it in relation to the lipid bilayer surface (Fig. 1a). This allowed the identification of 32 amino acid residues in βarr1 that potentially interact with lipid head groups, although ordered density for lipids was not observed. Two regions of arrestins had previously been suggested as interacting with lipids, the ‘344-loop’ (s18s19 loop) and the ‘197-loop’ (s11s12 loop)^26,27^. The nomenclature in parentheses is that implemented for arrestins in GPCRdb^28^. Both these loops in βarr1 contain backbone and side chains apparently buried in the head group region of the lipid bilayer and are flanked by residues where only the side chain interacts (Extended Data Figs. 8 and 9). Ten residues at the tip of the s18s19 loop were disordered and their position could therefore not be determined. Both the s11s12 and s18s19 loops in βarr1 are distant from the receptor in the complex. In contrast, two other regions of βarr1 that also appeared to interact with lipids (s8s9 and s15s16) contained residues that also interact with β_1_AR. The mutation L68C at the base of the finger loop is also accessible to the lipid bilayer, and is consistent with monobromobimane labelling studies, which shows a change in fluorescence when labelled arrestin couples to a GPCR^29^. Finally, the β-sheet s15 and the loop s14s15 contain many residues that could also potentially interact with lipids. It is notable that of the 18 residues in βarr1 that might interact with lipids through their side chains, 10 are either Lys or Arg. This suggests that negatively charged lipids such as phosphatidylinositols and/or phosphatidylserine, may play an important role in arrestin coupling^30^ *in vivo*, as has been suggested for coupling of the G protein G_s_^31^.

The structure of the formoterol-bound β_1_AR-βarr1 complex was compared with the β_2_AR-G_s_ complex^7^. The amino acid sequences of β_1_AR and β_2_AR are 59% identical (excluding the N-terminus and C-terminus) and have very similar inactive state structures (rmsds 0.4-0.6 Å)^17,19^ and active state structures coupled to nanobodies (rmsds 0.4-0.6 Å)^6,21,22^ so the comparison is reasonable. Superposition of β_1_AR and β_2_AR from the respective complexes (rmsd 1.0 Å, 1634 atoms) shows that the long axis of βarr1 is at ∼90° angle to the long axis of G_s_ (Extended Data Fig. 6). This could have an influence on the coupling efficiency of G proteins compared to arrestins if the GPCR form dimers^32^ and whether coupling occurs or not could be dictated by which transmembrane helices form the interface. The only significant difference in secondary structure between the different receptors is that the cytoplasmic end of H6 is an additional 7 Å away from the receptor in β_2_AR coupled to G_s_ compared to β_1_AR coupled to βarr (Fig. 2). The cleft in the intracellular face is thus 8 Å narrower when βarr1 is coupled to β_1_AR compared to G_s_ coupled to β_2_AR (measured between the Cα of Ser346-Arg284 in β_1_AR and the Cα of Ser329-Lys267 in β_2_AR). The amino acid residues forming the interface between β_1_AR and βarr1 are very similar to those forming the interface between β_2_AR and G_s_ (Fig. 3a). In particular, both complexes rely on extensive contacts between ICL2 and the cytoplasmic end of H3 of the receptor and either βarr1 or G_s_. However, there are contacts between the cytoplasmic ends of H2, H3, H7 and H8 in β_1_AR and βarr1 that are absent in the β_2_AR-G_s_ complex. There are also more extensive contacts between H5 and H6 of β_2_AR to G_s_ compared to the β_1_AR-βarr1 complex. The amino acid side chains in β_1_AR and β_2_AR at the respective interfaces are also similar in position, with the exception of Arg^3.50^ (Arg139 in β_1_AR and Arg131 in β_2_AR) in the DRY motif. In the β_2_AR-G_s_ complex and also in related complexes such as between the adenosine A_2A_ receptor and G_s_^33^, Arg^3.50^ extends away from the helix axis of H3 to form an interface between Tyr391 of the G protein and the hydrophobic interior of the receptor (Fig. 3c). In contrast, Arg^3.50^ in the β_1_AR-βarr1 complex adopts a different rotamer making extensive polar interactions with Asp138^3.51^ and Thr76^2.39^ in the receptor and with Asp69 of βarr1 in the finger loop (Fig. 3b). The rotamer of Arg^3.50^ and its interactions to other β_1_AR side chains in the β_1_AR-βarr1 complex is virtually identical to those observed in inactive state structures of β_1_AR and in active state structures stabilised by nanobodies^17,21^. One final observation in the comparison between the β_1_AR-βarr1 complex and the β_2_AR-G_s_ structure is that the α5 helix of G_s_ does not overlap precisely with the position of the finger loop of βarr1 in the β_1_AR-βarr1 complex, with the finger loop lying closer to H7-H8 than the α5-helix (Figs. 3d & 3e).

**Fig. 3.**
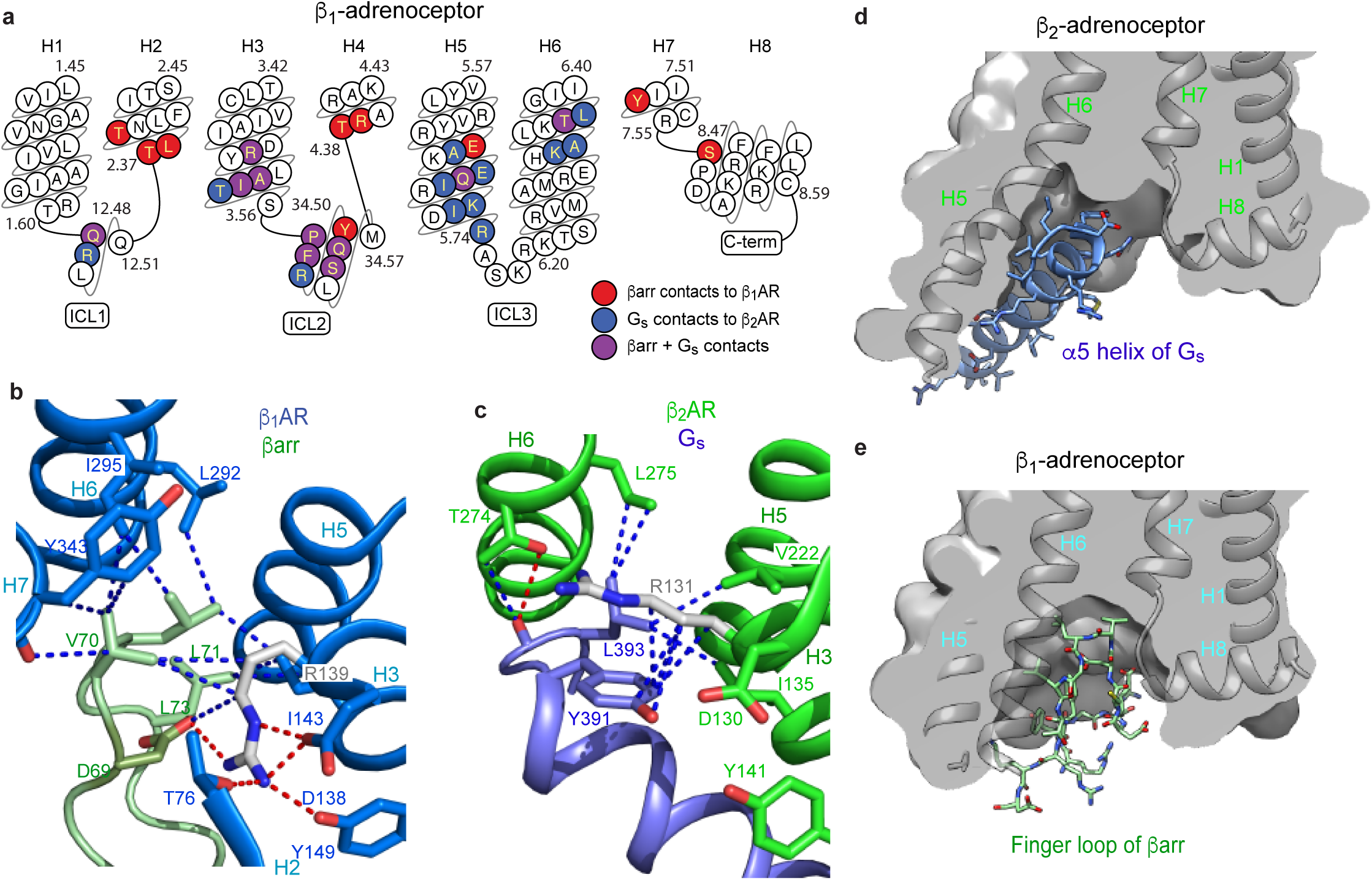
Comparison of the receptor coupling interfaces in the complexes of β_1_AR-βarr1 and β_2_AR-G_s_. **a,** snake plot of the intracellular region of turkey β_1_AR with amino acid residues colour coded according to interactions: red, contact between β_1_AR and βarr1; blue, contact between β_2_AR and G_s_; purple, both of the previously mentioned contacts. **b**, detail of the interface between the βarr1 finger loop (pale green) and β_1_AR (pale blue). **c**, detail of the interface between the α5 helix of the G_s_ α-subunit (blue) and β_2_AR (green). In panels **b** and **c**, depicted are polar interactions (red dashes), Van der Waals interactions (blue dashes; atoms ≤ 3.9 Å apart) and Arg^3.50^ (sticks; carbon, grey). **d**-**e**, cross-sections through the intracellular halves of β_1_AR and β_2_AR to highlight the different shapes of the intracellular cleft formed upon coupling of βarr1 compared to G_s_. Transmembrane helices are shown for orientation and they are in front of the cross-section.

Experimental data suggest that formoterol, carmoterol and carvedilol are arrestin biased ligands and that isoprenaline is a balanced agonist signalling equally between the G protein and arrestin pathways^5,34^ (see Fig. 4d for ligand structures). To identify elements that may be involved in biased signalling we therefore compared formoterol-bound β_1_AR coupled to βarr1 and to Nb80 (Nb80 being a known mimetic of the G protein G_s_^6^). The largest differences were observed on the intracellular face of β_1_AR where the ends of H5 and H6 were closer to the receptor core by 6.7 Å (Cα of Ile241) and 1.9 Å (Cα of Lys284), respectively, when βarr1 was coupled compared to when Nb80 was coupled (Fig 4b). On the extracellular face of the receptor (Fig 4c), the largest change is in ECL3 that occludes the entrance to the orthosteric binding pocket through a 2.2 Å shift in its position (as measured at Cα of Arg317). There was no significant density for side chains in ECL3 of the β_1_AR-βarr1 structure, so we cannot compare changes in their interactions. In the orthosteric binding site there was a significant difference in the interactions between formoterol and the residues Ser211^5.41^ and Ser215^5.46^ (Fig. 4a). Three potential hydrogen bonds between formoterol and either Ser215^5.46^ (one hydrogen bond) or Ser211^5.41^ (two hydrogen bonds) are reduced in length by 0.4 Å, 0.8 Å and 0.2 Å, respectively, which is consistent with the decreased affinity of formoterol when βarr1 is coupled compared to mini-G_s_ (Extended Data Fig. 1). This also correlates with the inward movement of the cytoplasmic end of H5 by 6.7 Å. In comparison to isoprenaline, formoterol also makes extra contacts to both H6 (Val326^7.36^ and Phe325^7.35^) and ECL2 (D200), similar to the additional contacts observed in β_1_AR structures with bound carmoterol^17^ or carvedilol^23^ (Fig. 4a). Previous work identified a causal link between G protein coupling and changes in the extracellular positions of H6-ECL3-H7 that resulted in the decreased rates of ligand association/dissociation^35^ and an increase in ligand affinity^21^. The β_1_AR-βarr1 structure suggests that this region is also implicated in ligand bias, with the arrestin biased ligands of β-adrenoceptors possessing moieties that interact with the extracellular end of H6 and ECL2. This would be anticipated to affect the dynamics of the extracellular region which, in conjunction with the increased length of hydrogen bonds to H5, would affect the structure on the intracellular face where G proteins and arrestins couple.

**Fig. 4.**
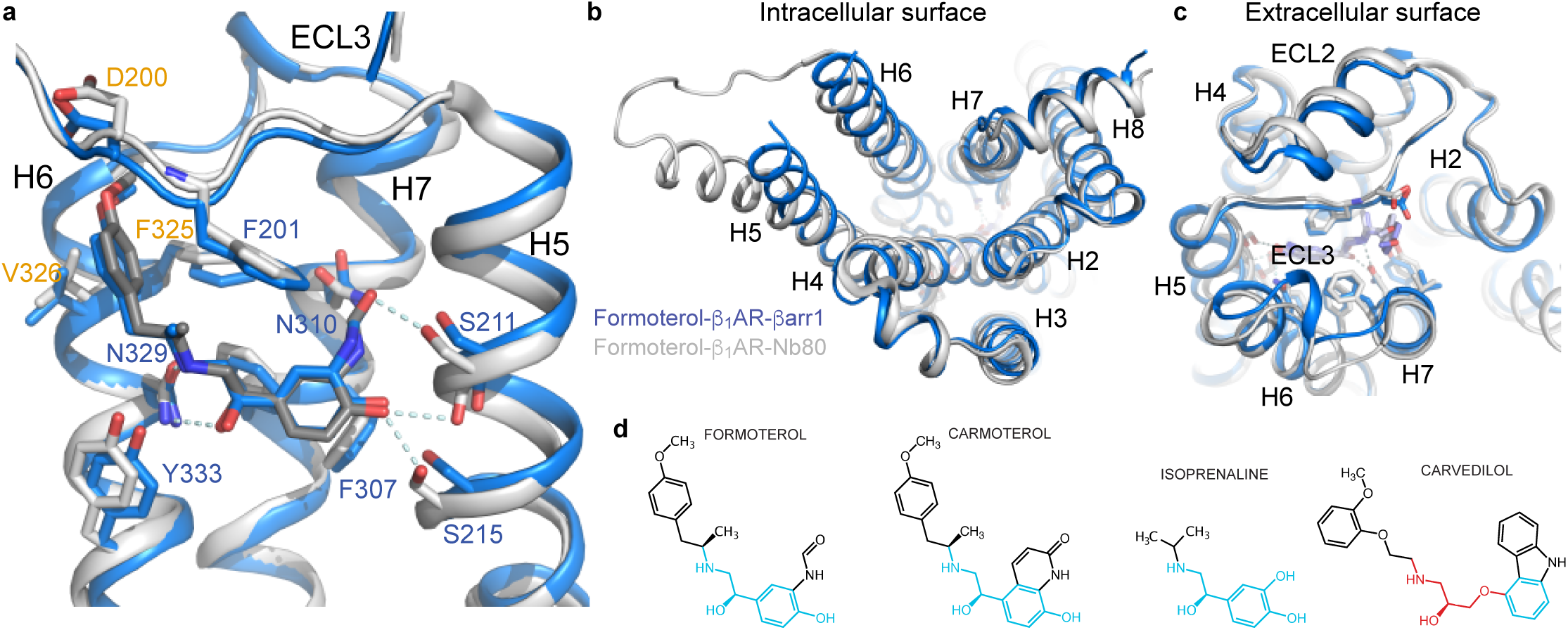
Differences between formoterol-bound β_1_AR coupled to either βarr1 or Nb80. **a-c**, superposition of β_1_AR coupled to βarr1 (blue cartoon) and β_1_AR coupled to Nb80 (grey cartoon) with residues interacting with the ligand shown as sticks. Residues labelled in orange interact with formoterol but not isoprenaline. **d**, structures of arrestin biased ligands (formoterol, carmoterol, carvedilol) and a balanced agonist (isoprenaline). Regions in blue are identical to adrenaline and the red region in carvedilol is the oxypropylene linker typical of β-adrenoceptor antagonists of the G protein pathway.

The structure of β_1_AR-βarr1 suggests possibilities of designing biased agonists that result in repositioning of H5 compared to balanced agonists, and also new opportunities for developing drugs that bind on the intracellular surface of receptors. The difference in position of the α5 helix in β_2_AR-G_s_ away from H7-H8 offers the possibility of a small molecule drug binding in this region that would not affect G_s_ coupling, but would sterically block βarr1 binding, resulting in G protein-biased signalling. Targeting the interfaces between H6 and other helices to prevent the larger displacement of H6 in the β_2_AR-G_s_ structure compared to the β_1_AR-βarr1 structure would result in arrestin-biased signalling. These approaches are plausible given that allosteric ligands are known to interfere with receptor signalling when bound to the intracellular region of a receptor^36^ whilst others bind to the lipid-exposed surface of transmembrane domains^37^. The challenge remains in designing compounds that specifically target these sites.

## Acknowledgements

The work in C.G.T.’s laboratory was funded by a grant from the European Research Council (EMPSI 339995), Heptares Therapeutics Ltd and core funding from the Medical Research Council [MRC U105197215]. The research program in A.K.S’s laboratory is supported by an Intermediate Fellowship of the Wellcome Trust/DBT India Alliance Fellowship (grant number IA/I/14/1/501285), the Science and Engineering Research Board (SERB) (EMR/2017/003804), Innovative Young Biotechnologist Award from the Department of Biotechnology (DBT) (BT/08/IYBA/2014-3) and the Indian Institute of Technology, Kanpur. H.D.-A. is supported by the National Post-Doctoral Fellowship of SERB (<u>PDF/2016/002930</u>) and DBT-BioCaRE grant (BT/PR31791/BIC/101/1228/2019). We thank Diamond Light Source (UK) for access and support of the cryo-EM facilities at eBIC (proposal EM17434) funded by the Wellcome Trust, MRC and BBSRC. We thank the beamline staff at the European Synchrotron Radiation Facility MASSIF-1 for help with X-ray diffraction data collection. We thank T. Nakane and P. Kolb for helpful discussions, D. Gloriam for access to unreleased data from GPCRdb, and G. Cannone from the LMB EM facility and J. Grimmett and T. Darling from LMB scientific computing for technical support during this work.

## Author contributions

Y.L. performed receptor, arrestin and zap1 expression, purification, nanodisc reconstitution and complex formation, cryo-EM grid preparation, data collection, data processing and model building. T.W. performed receptor and nanobody expression, purification and complex formation, crystallization, cryo-cooling of the crystals, X-ray data collection, data processing, and X-ray structure refinement. Y.L. and T.W. performed the pharmacological analyses. S.P. and H.D.-A. performed F_ab_ expression, purification and validation. R.N. developed the sortase ligation of phosphorylated peptides onto β_1_AR. P.C.E purified mini-G_S_. J.G.-N. advised on cryo-EM data collection, data processing and model building. A.G.W.L. advised on X-ray data processing, structure solution and analysis. Y.L. and C.G.T. carried out structure analysis and manuscript preparation. A.K.S. managed the production of F_ab_. C.G.T. analysed data and managed the overall project. The manuscript was written by C.G.T and Y.L., and included contributions from all the authors.

## Author information

Reprints and permissions information is available at www.nature.com/reprints. The authors declare the following competing interests: CGT is a shareholder, consultant and member of the Scientific Advisory Board of Heptares Therapeutics, who also partly funded this work. Correspondence and requests for materials should be addressed to.

## Materials and Methods

### Cloning, expression and purification of β_1_AR

The turkey (*Meleagris gallopavo*) β_1_AR constructs used for crystallization (see Extended Data Fig. 10) of the β_1_AR-Nb80 complex (trx-β_1_AR) was based on β44-m23^17^. The construction of trx-β_1_AR has been described previously^21^. Relative to wild-type β_1_AR, trx-β_1_AR contains truncations at the N- and C-termini (upstream of A33 and downstream of L367, respectively), and in intracellular loop 3 (C244 to R271, inclusive). Thioredoxin (*E. coli* trxA, with mutations C32S & C35S) was attached to the N-terminus via the linker EAAAK. trx-β_1_AR also contains the four thermostabilising mutations (R68S^1.59^, M90V^2.53^, F327A^7.37^ and F338M^7.48^), as well as two additional mutations C116L^3.27^ and C358A^8.59^. A hexahistidine tag is fused to the C-terminus of trx-β_1_AR.

The turkey β_1_AR construct used for electron cryo-microscopy of the β_1_AR-βarr1-F_ab_30 complex (β83S) was also based on β44-m23^17^. β83S (see Extended Data Fig. 10) shares the same truncations at the N-terminus and in intracellular loop 3 as trx-β_1_AR. β83S contains six thermostabilising mutations (M44C^1.35^, M90V^2.53^, V103C^2.66^, D322K^7.32^, F327A^7.37^ and F338M^7.48^), as well as three additional mutations C116L^3.27^, E130W^3.41^ and C358A^8.59^. The sequence downstream of C358A^8.59^ has been replaced with a linker sequence mimicking the C-terminal tail of vasopressin receptor 2 (V_2_R). The sequence contains a sortase recognition site (LPETG) followed by a heptahistidine tag [ARGRPLPETGGGRRHHHHHHH]. The sortase site is positioned in order to maintain the relative distance between H8 in V_2_R and the conserved phosphoserine triad motif following sortase assembly (Extended Data Fig. 4). β84S is identical to β83S except for an N-terminal MBP domain fusion constructed with a flexible linker.

The generation of trx-β_1_AR baculovirus and its expression and subsequent purification was performed as described previously^21^. It was solubilized and purified in decylmaltoside (DM, Generon) and eluted off the alprenolol sepharose ligand affinity column as described previously^17,38,39^ with 100 µM formoterol. The β83S construct was cloned into the baculovirus transfer vector pBacPAK8 (Clontech). Baculovirus expressing β83S was prepared using the flashBAC ULTRA system (Oxford Expression Technologies Ltd). β83S was expressed in *Trichoplusia ni* cells (Expressions Systems). Cells were grown in suspension in ESF 921 medium (Expressions Systems) to a density of 3 x 10^6^ cells/ml. Cultures were supplemented with 5% BSA prior to infection with β83S baculovirus and incubated for 40 h. β83S was solubilized in 2% dodecylmaltoside (DDM, Generon) from the membrane fraction and further purified in 0.02% DDM by Ni^2+^-affinity chromatography and alprenolol sepharose ligand affinity chromatography. It was eluted from the alprenolol sepharose column with 100 µM alprenolol. β83S was further polished on a Superdex 200 Increase column to remove excess alprenolol.

β83_6P_ was generated by sortase A-mediated covalent assembly^40^ of purified β83S with a synthetic phosphopeptide, V_2_R_6P_ (GGGDE[pS]A[pT][pT]A[pS][pS][pS]LAKDTSS, Tufts University Core Facility). The expression plasmid for sortase A(P94S, D160N, D165A, K196T) was a gift from S. Eustermann and D. Neuhaus. β83S (1 mg/ml) was incubated overnight on ice with 10-fold molar excess of V_2_R_6P_ and His-tagged sortase A at 1:10 (mol/mol) enzyme:receptor ratio. The assembly mixture was pre-adjusted with NaOH to pH 7.5 prior to the addition of receptor. Unreacted receptor and enzyme were removed by negative Ni^2+^-affinity chromatography. β83_6P_ was further polished on a Superdex 200 Increase column.

### Expression and purification of nanobody Nb80

A synthetic gene (Integrated DNA Technologies) for Nb80^6^ was cloned into plasmid pET-26b(+) (Novagen) with a N-terminal His_6_ tag followed by a thrombin protease cleavage site. Expression in *E. coli* strain BL21(DE3)RIL (Agilent Technologies) and purification from the periplasmic fraction was as described elsewhere^22^, but with removal of the His_6_ tag was by a thrombin (Sigma) protease cleavage step before concentration to 40 mg/ml.

### Formation of formoterol-bound trx-β_1_AR-nanobody complex and purification with detergent exchange by size exclusion chromatography

Preparation of receptor-nanobody complex was performed as described previously^21^. Formoterol-bound trx-β_1_AR (1.5 mg) was mixed with 1.5-fold molar excess nanobody (0.65 mg), cholesteryl hemisuccinate (Sigma) was added to 0.1 mg/ml in a final volume of 150 µL, and then incubated for 2 h at room temperature.

After incubation, size exclusion chromatography (SEC) was performed to separate receptor-nanobody complex from excess nanobody and to exchange the detergent from DM to HEGA-10 (Anatrace) for crystallization by vapour diffusion. A Superdex 200 10/300 GL Increase column (GE Healthcare) was used at 4 °C, the column was equilibrated with SEC buffer (10 mM Tris-Cl^-^ pH 7.4, 100 mM NaCl, 0.1 mM EDTA, 0.35% HEGA-10) supplemented with 10 µM formoterol. Samples containing complex were mixed with 200 µl SEC buffer and centrifuged (14,000 x *g*, 5 minutes) immediately prior to SEC (flow rate 0.2 ml/minute), with a run time of one hour which was sufficient for a near-complete detergent exchange as indicated by quantitation of residual glycosidic detergent^41^. Peak fractions corresponding to complex were concentrated to 15 mg/ml for crystallization by vapour diffusion using Amicon Ultra-4 50 kDa centrifugal filter units (EMD-Millipore).

### Crystallization of receptor-nanobody complex, data collection, processing and refinement

Crystals were grown in 150 + 150 nl sitting drops by vapour diffusion at 18 °C against reservoir solutions containing 0.1 M HEPES-Na^+^ pH 7.5 and 21-24% PEG1500. Crystals usually appeared within 2 h and grew to full size (up to 200 µm in length) within 48 h. Crystallization plates were equilibrated to 4 °C for at least 24 hours before cryo-cooling. Crystals were picked with LithoLoops (Molecular Dimensions Ltd) and dipped in 0.1 M HEPES-Na^+^ pH 7.5, 25% PEG1500, 5% glycerol before plunging into liquid nitrogen.

Diffraction data for the trx-β_1_AR-Nb80 complex with formoterol bound were collected at ESRF, Grenoble using the autonomous beamline MASSIF-1^42^. X-ray diffraction data were collected from a single point on the crystal using automatic protocols for the location and optimal centring of crystals^43^. The beam diameter was selected automatically to match the crystal volume of highest homogeneous quality and was therefore collimated to 30 µm, and strategy calculations accounted for flux and crystal volume in the parameter prediction for complete data sets^44^. Diffraction data were processed using MOSFLM^45^ and AIMLESS^46^, the structure was solved using PHASER^47^ with use of the crystal structures of the active state β_2_AR stabilized with nanobody Nb80^6^ and wild-type thioredoxin (PDB codes 3P0G, 2H6X) as search models. Diffraction was anisotropic, as indicated by the estimated resolution limits (CC_1/2_=0.3) in h,k,l directions (Extended Data Table 2). In order to retain statistically significant diffraction data, but eliminating reflections in less well diffracting directions, the data were truncated anisotropically using the UCLA Diffraction Anisotropy Server (http://services.mbi.ucla.edu/anisoscale/). Model refinement and rebuilding were carried out with REFMAC5^48^ and COOT^49^.

Cloning, expression and purification of human β-arrestin-1. **Wild-type human** βarr1 was cloned into the pTrcHisB vector with a TEV protease-cleavable N-terminal His_6_ and FLAG tag. Two mutations were introduced by site-directed mutagenesis: L68C, a finger loop-mutation commonly used in the functional labelling of arrestins^29^, and R169E, which disrupts the polar core and predisposes arrestin to activation^50^. Arrestin was expressed in BL21 cells. Cells were grown in LB medium supplemented with 100 µg/ml ampicillin at 25 °C. Expression was induced with 30 µM IPTG at a cell density of OD_600_ 0.5. The temperature was lowered to 15 °C and the cells allowed to grow for an additional 20 h. Cells were harvested and flash frozen in liquid nitrogen and stored at –80 °C. Arrestin was purified sequentially by Ni^2+^-affinity chromatography, TEV protease-cleavage of its N-terminal affinity tags, and Heparin chromatography, eluting off the Heparin column using 1 M NaCl. Purified arrestin was further polished on a Superdex 200 prep grade column (GE Healthcare) equilibrated in 20 mM Tris-Cl^-^ pH 8.0, 0.1 M NaCl, 10% glycerol, 0.5 mM DTT. Peak fractions were pooled and concentrated to 20 mg/ml and flash frozen as aliquots in liquid nitrogen and stored at –80 °C.

### Expression and purification of zebra fish apo-lipoprotein A-1

Zebra fish apo-lipoprotein A-1 (zap1) was expressed and purified as previously described^51^. Briefly, a pET-28a vector harbouring zap1 with a HRV-3C protease-cleavage N-terminal His_6_ tag was transformed into *E. coli* BL21(DE3)RIL cells. Cells were grown at 37°C in TB medium supplemented with kanamycin. Expression was induced at OD_600_ 1-1.5 with 1 mM IPTG. The temperature was lowered to 25°C and the culture was allowed to grow for 3 h. Cells were lysed by sonication in the presence of 1% (v/v) Triton X-100. Cell lysate was clarified by centrifugation and passage through a 0.22µm filter prior to loading onto a HisTrap-FF column. The pellet from the previous centrifugation step was resuspended in buffer containing 6 M guanidine hydrochloride (GnHCl), clarified by centrifugation, and loaded onto the HisTrap column. The column was washed in successive buffers (base: 20 mM Tris-Cl^-^ pH 7.5, 0.3 M NaCl, 20 mM imidazole) containing first, 6 M GnHCl, then 0.2% Triton X-100, followed by 50 mM Na-cholate, before eluting in 20 mM Tris-Cl^-^ pH 7.5, 150 mM NaCl, 20 mM Na-cholate, 0.3 M imidazole. Purified zap1 was treated with HRV-3C protease in the presence of 0.5 mM TCEP to remove the His_6_ tag prior to polishing on a Superdex 200 Increase column equilibrated in 20 mM Tris-Cl^-^ pH 7.5, 150 mM NaCl, 20 mM Na-cholate.

### Expression and purification of F_ab_30

The coding region of F_ab_30 was synthesized by GenScript based on previously published crystal structure (PDB 4JQI). For large-scale purification, F_ab_30 was expressed in the periplasmic fraction of *E. coli* 55244 cells (ATCC) and purified using Protein L (GE Healthcare) gravity flow affinity chromatography as published previously^29^. Briefly, Cells harboring F_ab_30 plasmid were used to inoculate 2xYT and grown overnight at 30 °C. Fresh 2xYT was inoculated with 5% initial inoculum and grown for an additional 8 h at 30°C. Cells were harvested and resuspended in an equal volume of CRAP medium supplemented with ampicillin, and grown for 16 h at 30 °C.

For F_ab_30 purification, cells were lysed in Fab-lysis buffer (50 mM HEPES-Na^+^ pH 8.0, 0.5 M NaCl, 0.5% (v/v) Triton X-100, 0.5 mM MgCl_2_) by sonication.

Crude cell lysate was heated in a 65°C water bath for 30 min and chilled immediately on ice for 5 min. Subsequently, lysate was clarified by centrifugation at 20,000 x *g* and passaged through pre-equilibrated Protein L resin packed gravity flow affinity column. After binding at room temperature, beads were washed extensively with wash buffer (50 mM HEPES-Na^+^ pH 8.0, 0.5 M NaCl). Protein was eluted with 100 mM acetic acid into tubes containing 10% vol. neutralization buffer (1 M HEPES pH 8.0). Following elution, sample was desalted into Fab-storage buffer (20 mM HEPES pH 8.0, 0.1 M NaCl) using a pre-packed PD-10 column (GE Healthcare). Purified F_ab_30 was flash frozen stored in buffer supplemented with 10% glycerol.

### Functional validation of purified F_ab_30

Functionality of purified F_ab_30 was assessed using co-immunoprecipitation with their reactivity towards V_2_Rpp-bound βarr1 as readout following a previously published protocol^52^. Briefly, F_ab_30 (1.5 µg) was incubated with purified βarr1 (2.5 µg) in the presence or absence of V_2_Rpp (pre-incubated with 5-10 fold molar excess compared to βarr1) in 100-200µl reaction volume. After 1h incubation at room temperature, pre-equilibrated Protein L beads were added to the reaction mixture and incubated for an additional 1 h. Subsequently, Protein L beads were washed 3-5 times using 20 mM HEPES-Na^+^ pH 7.4, 150 mM NaCl, 0.01% MNG to remove any non-specific binding. Bound proteins were eluted using 2×SDS loading buffer and separated by SDS-PAGE. Interaction of F_ab_30 with activated βarr1 was visualized using Coomassie-staining and Western blot.

### Reconstitution of purified β_1_AR into nanodiscs and complex formation

Reconstitution was performed by adapting established protocols^53^. Stocks of 16:0-18:1 PC (POPC) and 16:0-18:1 PG (POPG, Avanti Polar Lipids) in chloroform were dried down under a nitrogen stream and fully solubilised in 20 mM HEPES-Na^+^, 150 mM NaCl, 100 mM Na-cholate to make 50 mM lipid stocks. β83_6P_ (500 µg) was reconstituted into zap1-supported nanodiscs containing 7:3 (mol/mol) POPC:POPG. Receptor, zap1 and lipids at a molar ratio of 1:10:1000 (net. 18 mM cholate) were mixed and incubated for an hour on ice. A three-fold excess of damp, pre-equilibrated Bio-Beads SM-2 (Bio-Rad) was added in batch and the sample was mixed end-over-end overnight at 4 °C. An absorption capacity of 80 mg cholate/g was used to calculate the requisite amount of polystyrene beads^54^. The reconstituted sample, composed of a mixture of β83_6P_-incorporated nanodiscs and zap1/lipid-only nanodiscs, was further polished on a Superdex 200 Increase column equilibrated in 20 mM HEPES-Na^+^, 150 mM NaCl, 5 µM formoterol.

The nanodisc mixture was supplemented with a further 10 µM formoterol and incubated with a 2-fold excess of βarr1(L68C, R169E) for 1 h on ice. A 2-fold excess of His-tagged F_ab_30 was added and the mixture incubated for 1 h. Subsequently, the mixture was left to incubate in batch with 1 mL Ni-NTA resin (QIAGEN) overnight at 4°C. A pull-down of β83_6P_-βarr1-F_ab_30 complex in nanodisc was performed by Ni^2+^-chromatography exploiting His-tagged F_ab_30 to remove tag-free zap1/lipid-only nanodiscs. The nanodisc-embedded ternary complex was separated from excess F_ab_30 on a Superdex 200 Increase column equilibrated in 10 mM HEPES-Na^+^ pH 7.5, 20 mM NaCl, 2 µM formoterol. SEC fractions were either used immediately for cryo-EM grid preparation or divided into aliquots and flash frozen and stored at –80°C. Grids prepared with freshly isolated complex or samples that had been subjected to a freeze/thaw cycle were identical in apparent quality.

### β_1_AR-β-arrestin1-F_ab_30 cryo-grid preparation and data collection

Cryo-EM grids were prepared by applying 3 µL sample (at a protein concentration of 1 mg/ml) on glow-discharged holey gold grids (Quantifoil Au 1.2/1.3 300 mesh). Excess sample was removed by blotting with filter paper for 2-3 s before plunge-freezing in liquid ethane (cooled to −181 °C) using a FEI Vitrobot Mark IV maintained at 100% relative humidity and 4 °C. Data collection was carried out on grids made from a single preparation of β_1_AR-βarr1-F_ab_30 complex. Images were collected on a FEI Titan Krios microscope at 300 kV using a GIF Quantum K2 (Gatan) in counting mode. Data were collected in 3 independent sessions—two on LMB-Krios2; one on Diamond eBIC-Krios1—to give a total of 18,581 movies. When processing previous datasets, particles were assessed by cryoEF^55^ to have an orientation distribution efficiency, E_od_ ∼0.55, indicating moderately severe preferential orientation of the particles in freestanding ice. In order to improve orientation distribution, micrographs in this study were collected with a 30°-stage tilt. On LMB-Krios2, automated data acquisition was performed using serialEM^56^. Grid squares were subdivided into 3×3 grid hole-matrices. Stage shift was used to align the central grid hole within the acquisition template. Subsequently, image shift with active beam-tilt compensation was used to record from the nine holes. Large changes in sample height due to stage-tilt were compensated for by an equivalent degree of defocus adjustment, pre-determined and applied so as to normalise to the target defocus value. On eBIC-Krios1, data collection was performed using EPU (Thermo Fisher Scientific). Stage shift was used to centre individual grid holes. In all sessions, two non-overlapping exposures, aligned along the tilt axis, were collected per grid hole. Micrographs were collected with a total accumulated dose of ∼45-50 e^-^/Å^2^. Each micrograph was collected as dose-fractionated movie frames (∼1.0 e^-^/Å^2^/frame) at a dose rate of 4.5 e^-^/pix/sec (LMB) or 3.3 e^-^/pix/sec (eBIC) with an energy selection slit width of 20 eV. The datasets were collected at a magnification of 105,000× (1.1 Å/pix, LMB) and 130,000× (1.047 Å/pix, eBIC).

### Data processing and model building

RELION-3.0.7 was used for all data processing unless otherwise specified^57^. Drift, beam-induced motion and dose-weighting were corrected in Warp-1.0.6 using a spatial resolution of 5×5 and a temporal resolution equal to the number of movie frames^58^. CTF estimation and determination of the focus gradient was performed in Warp using movie frame input, with 5×5 spatial resolution and a temporal resolution of 1. Micrographs were curated for quality based on ice contamination, CTF fitting quality, estimated resolution, and astigmatism, resulting in a trimmed dataset of 18,101 micrographs. Auto-picking was performed with a Gaussian blob as a template, which resulted in optimal particle picking. The CTF parameters for the picked coordinates were interpolated from the focus gradients modelled in Warp. Particles were extracted in a box-size equivalent to 264 Å and downscaled initially to 4.4 Å/pix. For each LMB-Krios2 session, micrographs were further separated into two halves, generating a total of five groups of particle stacks. Each group was processed independently. For each group, particles were subjected to two rounds of 3D classification in 6 classes using an *ab initio* model as reference. In the second round of 3D classification, particle distribution appeared to be dictated in part by the size of the nanodisc component (Extended Data Fig. 3). Aberrant classes of particles, such as C4 and C5 (Extended Data Fig. 3) were excluded from subsequent rounds of processing, as they probably arose from distorted nanodiscs (C4) or aggregation effects (C5) that arose during grid preparation. Particles of varying nanodisc sizes were combined, re-extracted with downscaling to 1.69 Å/pix, and refined to achieve an overall consensus alignment. Clear density could be observed for the transmembrane helices as well as two protrusions from the lipid boundary corresponding to ECL2 and ICL3, demarcating the volumes corresponding to receptor and zap1/lipid. Particles were subjected to Bayesian polishing before further refinement. Correcting for per-particle beam-induced motion consistently improved resolution by two resolution shells (according to a gold-standard FSC of 0.143). Signal subtraction was performed to remove most of the non-receptor component of the nanodisc, facilitating refinement of the β_1_AR-βarr1-F_ab_30 complex that included a thin annular layer of lipid. Subsequently, 3D classification without alignment into 6 classes (regularisation parameter, *T*=20) identified a subset of particles (∼8%) that refined to high resolution and showed fine map details in the receptor and arrestin regions. On trace-back, this subset of good particles constituted roughly an equal proportion of the class averages identified in the preceding round of 3D classification (*i.e.* class distributions based loosely on nanodisc morphology). At this stage, the good particles from the five groups were combined, re-extracted with downscaling of 1.1 Å/pix, and processed as a single dataset. The merged particle set was split according to microscope session for independent Bayesian polishing before re-merging for downstream processing. Following signal subtraction of the nanodisc and refinement, the model reached a resolution of 3.43 Å. Subsequently, refined particles were imported into and processed in RELION-3.1. On account of the image shift collection strategy used in LMB-Krios2, the particles from those two sessions were assigned to 1 of 18 optical groups—by sessions and based on position within their respective 3×3 matrices. Including the eBIC-Krios1 particles, this produced 19 optical group assignments, which were corrected independently for residual beam-tilt, anisotropic magnification, per-micrograph astigmatism, and per-particle CTF estimation. In the final refinement sequence, half maps were locally filtered between refinement iterations using SIDESPLITTER (K. Ramlaul, C.M. Palmer & C.H.S. Aylett, manuscript in preparation), an adaption of the LAFTER algorithm (Ramlaul *et al.* 2019) that maintains gold-standard separation between the two half maps. The final model contained 403,991 particles and reached an overall resolution of 3.3 Å with side chains visible for most of the complex (Extended Data Figs. 2 and 4). Local resolution estimates were calculated with RELION-3.1 showing the β_1_AR-βarr1 and βarr1-F_ab_30 interfaces at ∼3.2 Å and rising gradually to ∼3.7 Å at the level of the β_1_AR orthosteric binding site; H1 and the extracellular regions of the receptor, the C-distal end of arrestin, and CL-CH1 domains of F_ab_30 are at poorer resolution, with the worst regions reaching ∼4.5 Å at the most exposed edges. The final particle set was assessed to have an orientation distribution efficiency, E_od_ ∼0.72.

Model building and refinement was carried out using the CCP-EM^59^ and PHENIX^60^ software suites. The formoterol-bound trx-β_1_AR-Nb80 and βarr1-F_ab_30-V_2_Rpp crystal structures were used as starting models (PDB 6IBL and 4JQI). β_1_AR was modelled from A42 to A358 with a gap from R243 to R279 (inclusive) on account of weak density. The V_2_R_6P_ portion of β_1_AR has been modelled from E372 to D384. The intervening linker region to A358 was too flexible to be resolved. Density for all phosphoresidues was well resolved. βarr1 was modelled from T6 to E359, with a gap between R331 and S340 (inclusive), which constitutes a region encompassing the C-distal “344-loop” that potentially interacts with the lipid head group region. Initial manual model building was performed in COOT^49^ following a jelly-body refinement in REFMAC5^48^. Restraints for formoterol were generated using AM1 optimisation in eLBOW^61^. In order to better maintain geometry in the regions of weak density, secondary structure restraints, Ramachandran restraints and rotamer restraints were applied during real space refinement in PHENIX. The model followed iterative cycles of manual modification in COOT and restrained refinement in PHENIX. The final model achieved good geometry (Extended Data Table 1) with validation of the model performed in PHENIX, Molprobity^62^ and EMRinger^63^. The goodness of fit of the model to the map was carried out using PHENIX using a global model-vs-map FSC (Extended Data Fig. 4). Over fitting in refinement was monitored^64^ using FSC_work_/FSC_test_ by refining a ‘shaken’ model against half map-1 and calculating a FSC of the resulting refined model against half map-2.

### Expression and purification of mini-G_s_

Mini-G_s_ (construct R393) was expressed in *E. coli* strain BL21(DE3)RIL and purified by Ni^2+^-affinity chromatography, removal of the His tag using TEV protease and negative purification on Ni^2+^-NTA for TEV and undigested mini-G_s_ removal; and final SEC to remove aggregated protein^65^. Purified mini-G_s_ was concentrated to a final concentration of 100 mg/ml in 10 mM HEPES-Na^+^ pH 7.5, 100 mM NaCl, 10% v/v glycerol, 1 mM MgCl_2,_ 1 µM GDP and 0.1 mM TCEP.

### Radioligand binding studies on β_1_AR in nanodiscs

Purified β84S was inserted into nanodiscs and ligated to the phosphorylated peptide as described for β83S. Zap1 with lipid only (no receptor) was used to determine background binding. Nanodiscs containing either empty nanodiscs, β84_6P_, or β84_6P_ in the presence of βarr1 were prepared in the absence of ligands and then diluted into assay buffer for radioligand saturation binding studies as previously described for insect cell membranes^21^.

### Competition binding assays

Nanodiscs containing β84_6P_ were resuspended in 20 mM HEPES-Na^+^ pH 7.5, 50 mM NaCl, 2.5 mM MgCl_2_, 0.1% BSA. Aliquots were supplemented with mini-G_S_ construct R393 or βarr1 (final concentration 25 µM), either formoterol or isoprenaline (8 points, with the final concentration between 1 pM and 100 µM), and apyrase (final concentration 0.1 U/ml; only with mini-G_S_) to give a final volume of 120 µL or 220 µl. Samples were incubated at 20 °C for 1 h, before adding [^3^H]-DHA (Perkin Elmer) with concentrations of competing ligand in the range 1-2.5 x K_D_. Non-specific binding was determined by measuring binding in the presence of 100 µM unlabelled ligand. Samples were incubated at 20 °C for 1-5 h, before filtering through 96-well Multiscreen HTS GF/B filter plates (Merck Millipore) pre-soaked in 0.1% (w/v) polyethyleneimine, separating bound from unbound [^3^H]-DHA. Filters were washed three times with 200 µl chilled assay buffer, dried, and then punched into scintillation vials with 4 ml Ultima Gold scintillant (Perkin Elmer). Radioligand binding was quantified by scintillation counting with a Tri-Carb Liquid Scintillation Analyser (Perkin Elmer) and K_i_ values were determined using GraphPad Prism version 7.

---

**Extended Data Fig. 1.**
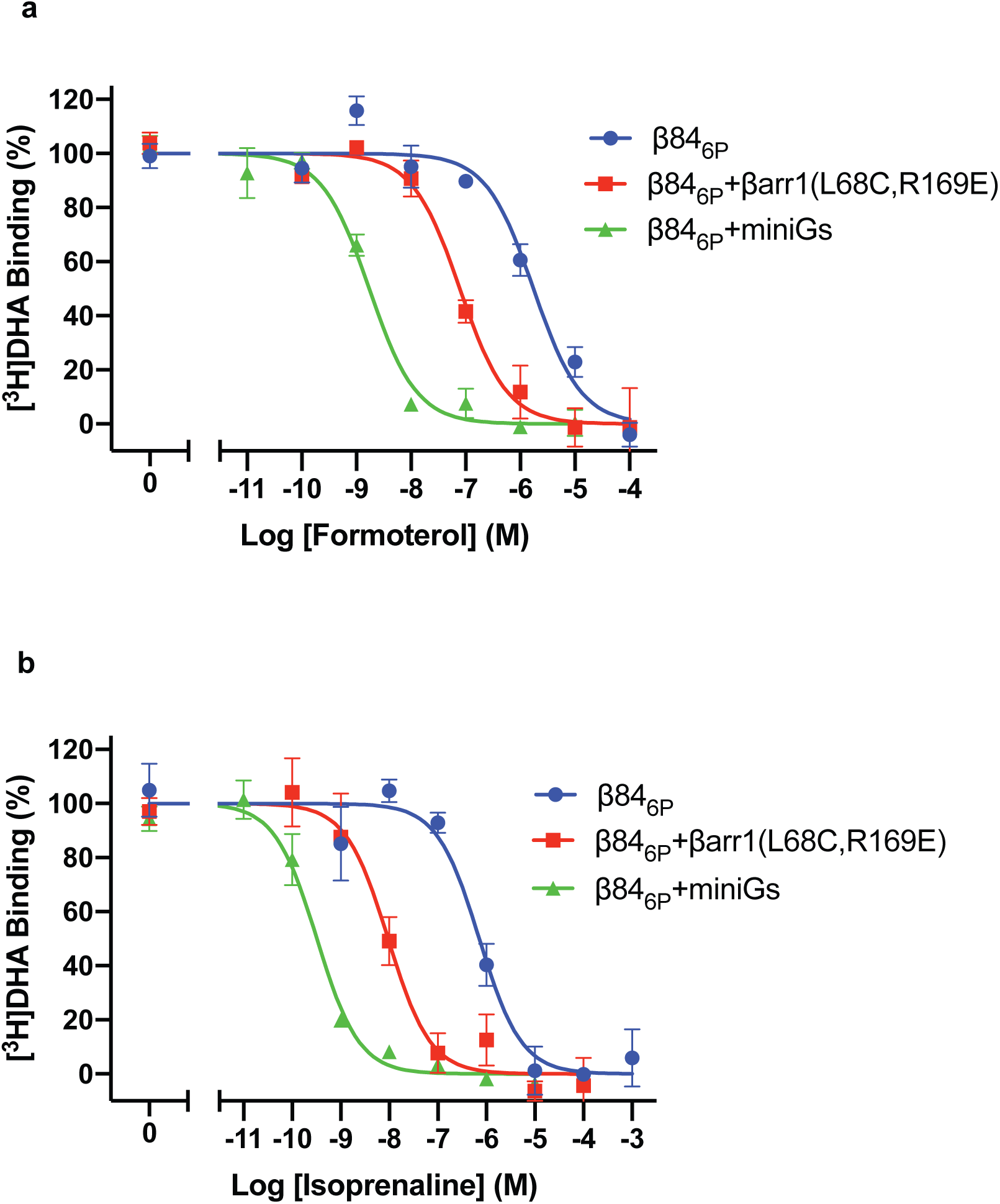
The modulation of β_1_AR agonist affinity by effector proteins. **a-b**, Representative competition binding curves using either formoterol or isoprenaline, respectively, show the high-affinity state of β_1_AR stabilised by either mini-G_s_ or βarr1. Experiments (see Methods) to determine the high affinity state were performed in a molar excess of mini-G_s_ (green curve) or βarr1 (red curve) and compared to the low affinity state (blue curves). Experiments were performed 2-4 times in duplicate and errors represent the SEM. The apparent K_i_s were determined using the Cheng-Prusoff equation and apparent K_d_s for _3_H-DHA of 6 nM (β84_6P_ and β84_6P_ + βarr1) and 1.5 nM (β84_6P_ + mini-G_s_). K_i_ values for formoterol are 1.5 ± 0.4 μM (β84_6P_), 42 ± 18 nM (β84_6P_ + βarr1) and 0.7 ± 0.1 nM (β84_6P_ + mini-G_s_). K_i_ values for isoprenaline are 340 ± 70 nM (β84_6P_), 4.4 ± 0.8 nM (β84_6P_ + βarr1) and 0.13 ± 0.02 nM (β84_6P_ + mini-G_s_).

**Extended Data Fig. 2.**
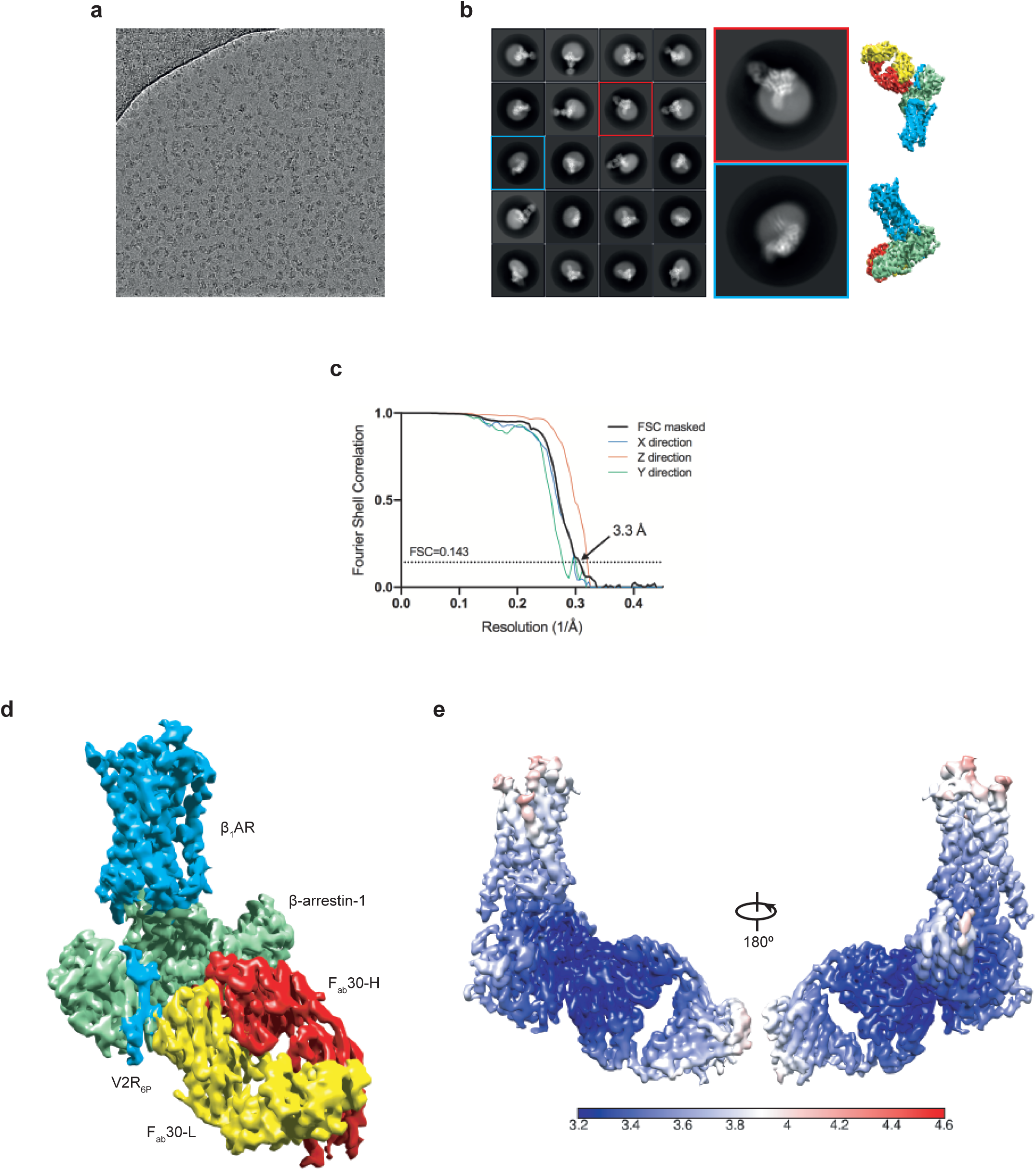
Cryo-EM single particle reconstruction of the β1AR-βarr1-Fab30 complex. **a,** Representative micrograph (magnification 105,000×, defocus –1.9 µm) of the β1AR-βarr1-Fab30 complex collected using a Titan Krios with the GIF Quantum K2 detector. **b,** Representative 2D class averages of the β1AR-βarr1-Fab30 complex determined using ∼1 million particles following 3D classification. Copies of the final reconstruction are juxtaposed to indicate relative orientations. **c,** FSC curve of the final reconstruction (black) showing an overall resolution of 3.3 Å using the gold standard FSC of 0.143. Shown in colour are the directional 3D-FSC curves calculated from the two half-maps66. **d,** Final reconstruction coloured by polypeptides (contour level 0.023). **e.** Local resolution estimation of the β1AR-βarr1-Fab30 map as calculated by RELION.

**Extended Data Fig. 3.**
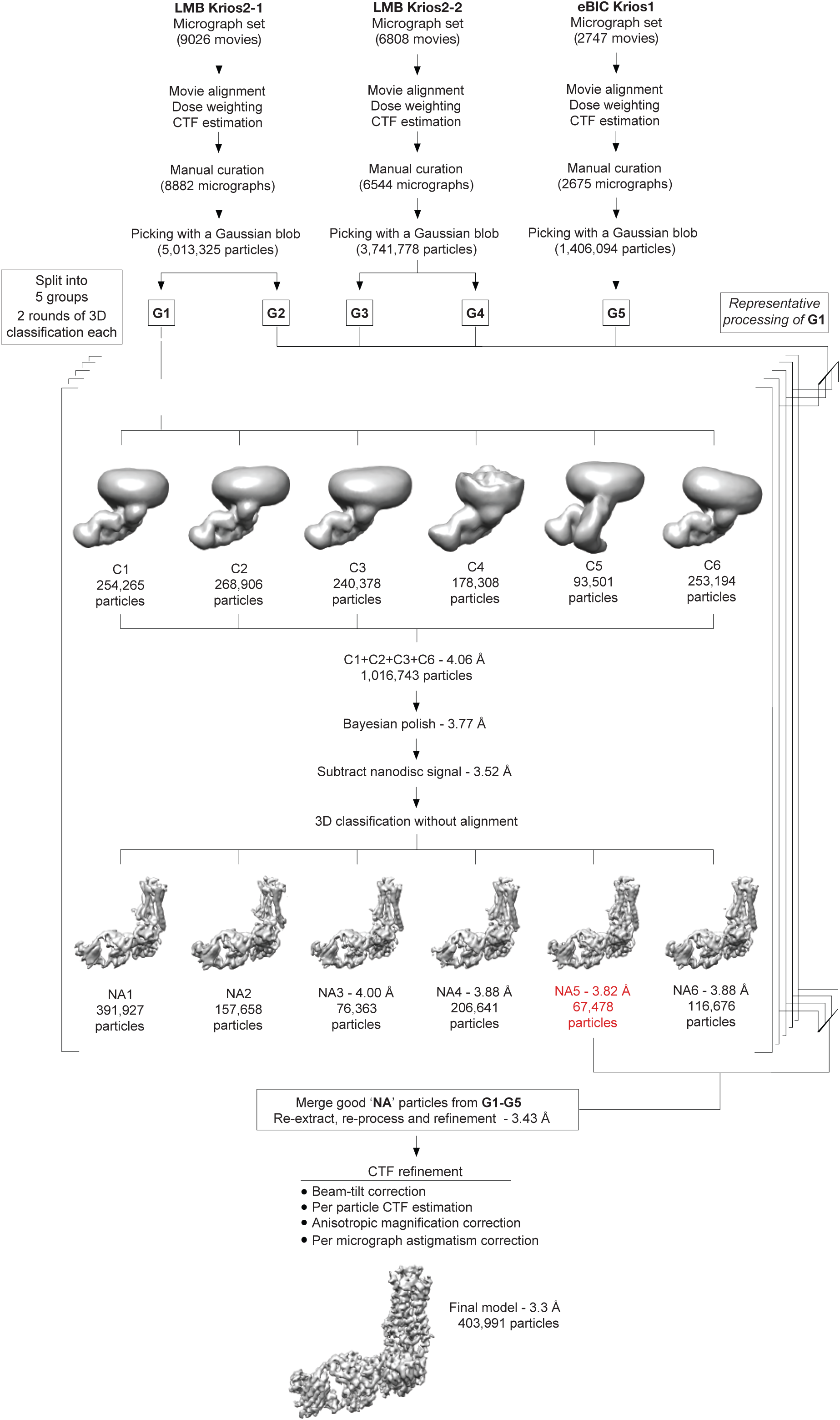
Flow chart of cryo-EM data processing. Micrographs were collected during three sessions on a Titan Krios (between 48 h and 96 h long) using a 30° stage tilt to reduce preferential particle orientation. Each dataset was corrected separately for drift, beam-induced motion and radiation damage. After focus gradient and CTF estimation, particles were picked using a Gaussian blob. At this stage, each of the LMB Krios2 datasets was split into 2 halves by micrographs, generating a total of five groups of particles. Each group was processed and curated independently. The number of particles from group G1 is indicated on the flowchart as a guide. At the bottom of the figure, the final number of particles is shown. Particles were submitted to two rounds of 3D classification using an *ab initio* model as a reference. In each round, classification was performed in six classes. The models with the best features were merged and refined together before correcting for per-particle beam-induced motion. Subtracted particles were generated by removing most of the non-receptor nanodisc signal and refined. 3D classification without alignment was performed in 6 classes using a mask encompassing the entire complex. The models showing the best features were refined either individually or in combination. The quality of the particles was judged based on both resolution and map features and weighed against the size of the particle set (the resolution of the models refers to the resolution after refinement and calculation of gold-standard FSC of 0.143). The best particles from each group were merged and re-extracted. Following merging, the combined particle set was processed together except at the stage of per-particle beam-induced motion correction, where particles were split into their session-stacks for Bayesian polishing. Following penultimate refinement, particles were assigned to one of 19 optical groups (see Methods) and corrected for beam-tilt, per-micrograph astigmatism, anisotropic magnification and per-particle CTF estimation. A final model with 403,991 particles was refined and achieved a global resolution of 3.3 Å.

**Extended Data Fig. 4.**
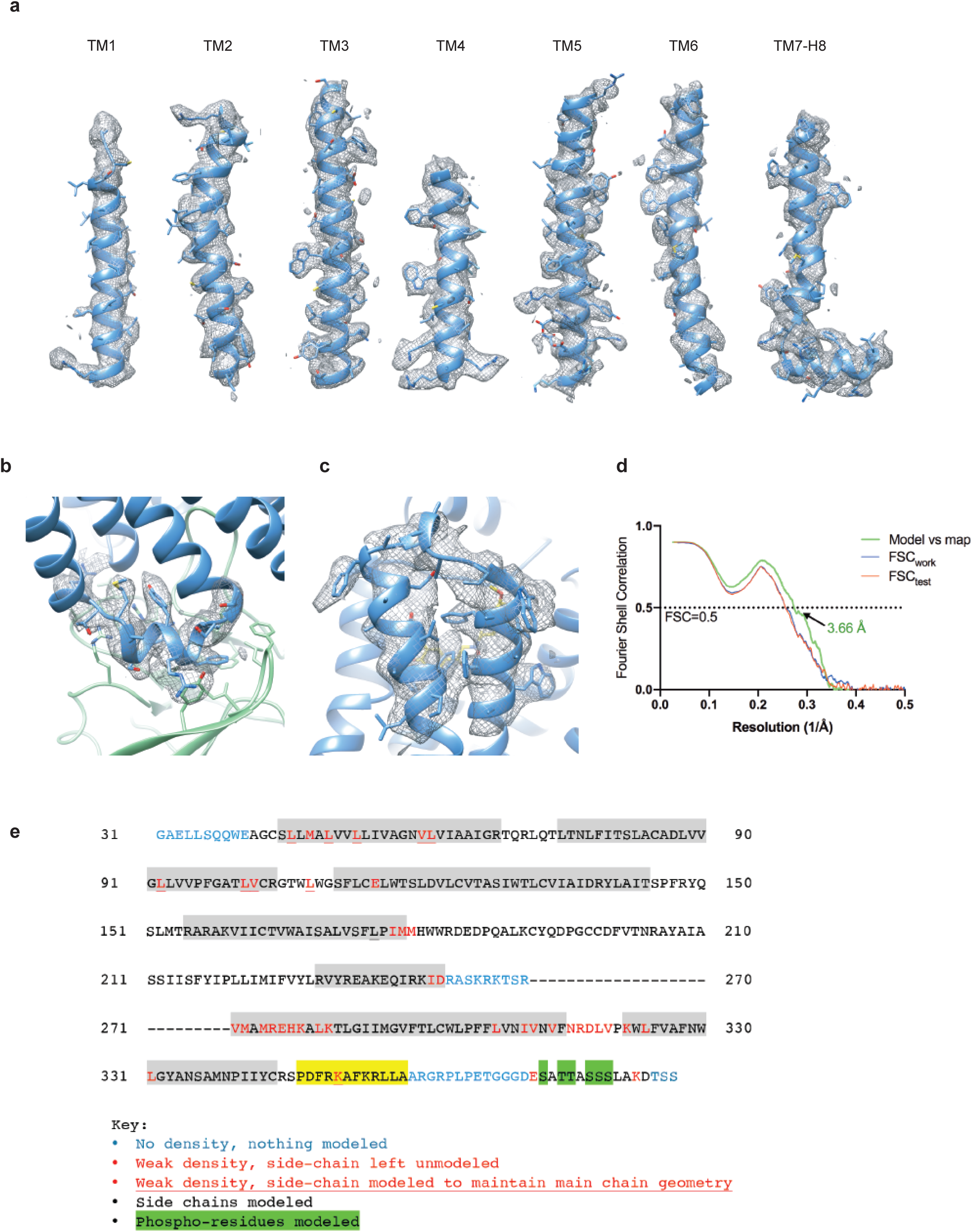
Cryo-EM map quality of the β_1_AR-βarr1-F_ab_30 complex and model validation. a, Transmembrane helices of β_1_AR with density shown as a mesh. **b,** Intracellular loop 2 of β_1_AR. For clarity, the neighbouring βarr1 side chains are depicted without density. **c,** Extracellular loop 3 of β_1_AR and the adjacent helical turns of H6 and H7. All density maps in panels **a**-**c** were visualised using Chimera (contour level 0.017) and encompass a radius of 2 Å around the region of interest. **d.** FSC of the refined model versus the map (green curve) and FSC_work_/FSC_free_ validation curves (blue and red curves, respectively). **e,** Amino acid sequence of the β_1_AR construct used for the cryo-EM structure determination. The residues are numbered according to the wild-type sequence of β_1_AR. Residues are coloured according to how they have been modelled. Black, good density allows the side chain to be modelled; red, limited density for the side chain, therefore the side chain has been truncated to Cβ; blue, no density observed and therefore the residue was not modelled. In some cases, side chains were included where there was only weak density as it aided maintenance of main chain geometry during restrained refinement. Regions highlighted in grey represent the transmembrane α-helices, amphipathic helix 8 is highlighted in yellow, and phosphorylated residues are highlighted in green. The dashes represent amino acid residues deleted.

**Extended Data Fig. 5.**
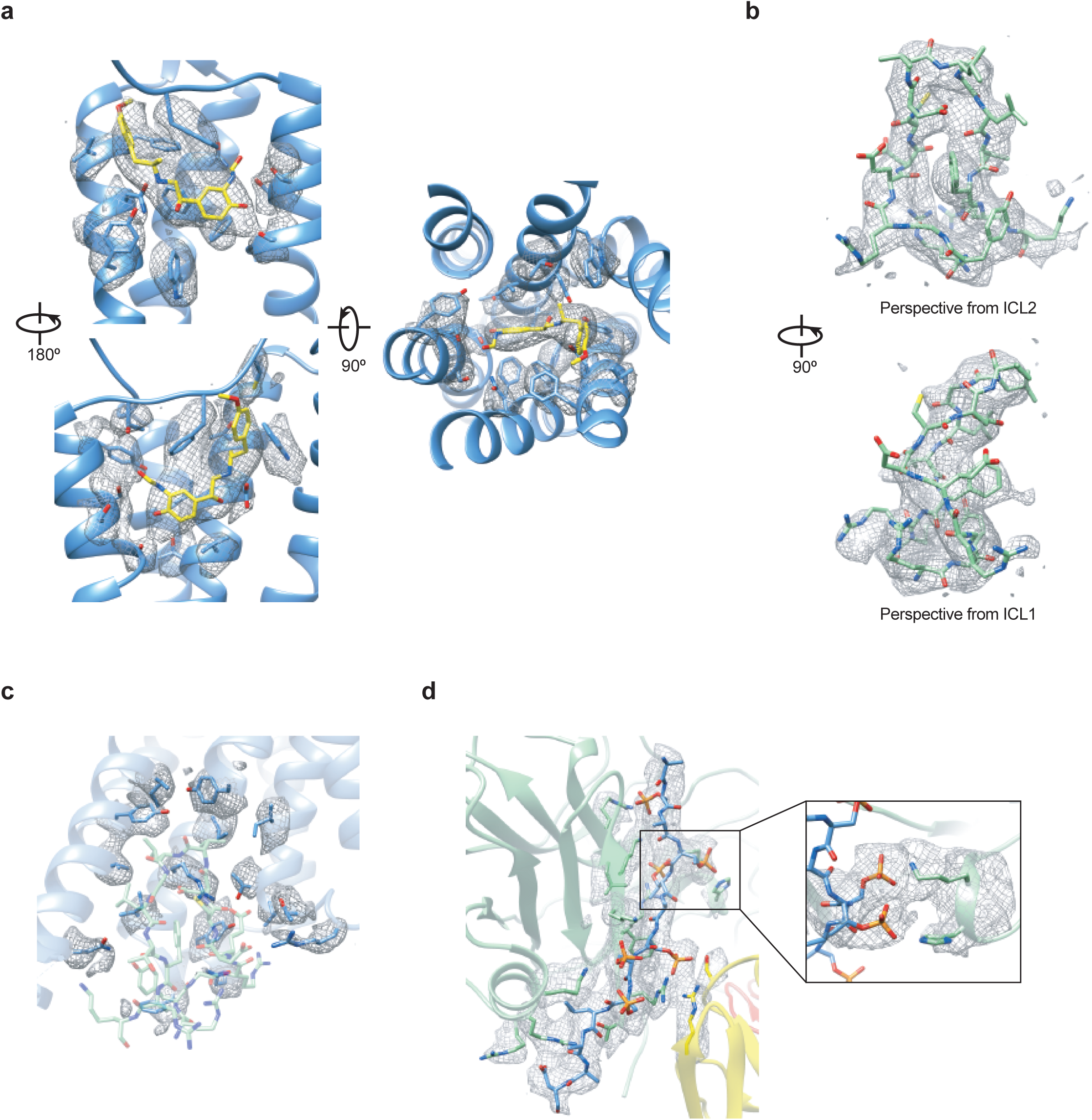
Cryo-EM map quality of β_1_AR-βarr1-F_ab_30 in the orthosteric binding site, arrestin-binding pocket and phosphorylated C-terminus. Unless otherwise stated, density maps (visualised in Chimera) are depicted with contour level 0.017 and encompass a radius of 2 Å around the region of interest. a, Formoterol and the neighbouring side chains in the orthosteric binding site. b, The finger loop of βarr1. c, The β_1_AR side chains neighbouring the finger loop of βarr1. d, The phosphorylated V_2_R_6P_ C-terminus. Inset, interaction between the V_2_R_6P_ phospho-threonine dyad and the βarr1 lariat loop. Density in the inset is depicted with contour level 0.01 (carve radius 2 Å).

**Extended Data Fig. 6.**
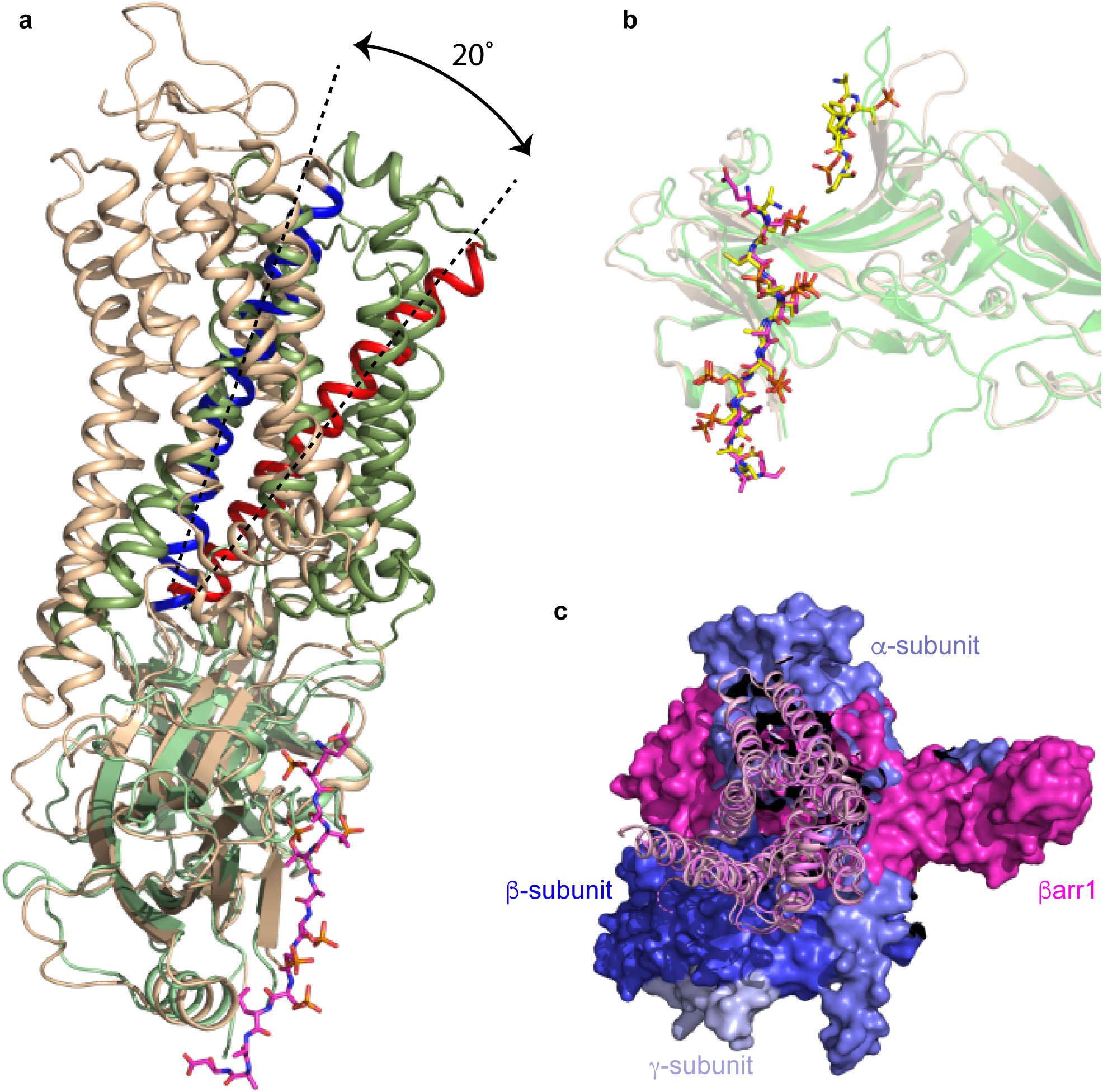
Comparison of the active states of arrestin. **a**, superposition of arrestin molecules in the complexes of β_1_AR-βarr1 (green) and rhodopsin-arrestin (pale brown). The different angle between the respective receptors and coupled arrestins is shown by the 20° difference in tilt of H3 (blue, H3 in rhodopsin; red, H3 in β_1_AR). **b**, superposition of the active state of βarr1 (pale brown; PDB code 4JQI) not bound to receptor and βarr1 (green) coupled to β_1_AR. The phosphopeptides are shown as sticks: yellow carbon atoms, V_2_Rpp in 4JQI; magenta carbon atoms, V_2_R_6P_ in the β_1_AR-βarr1 complex. **c**, superposition of β_1_AR and β_2_AR (pink and purple cartoons, respectively) coupled to either βarr1 (magenta surface) or G_s_ (blue and purple surfaces), respectively.

**Extended Data Fig. 7.**
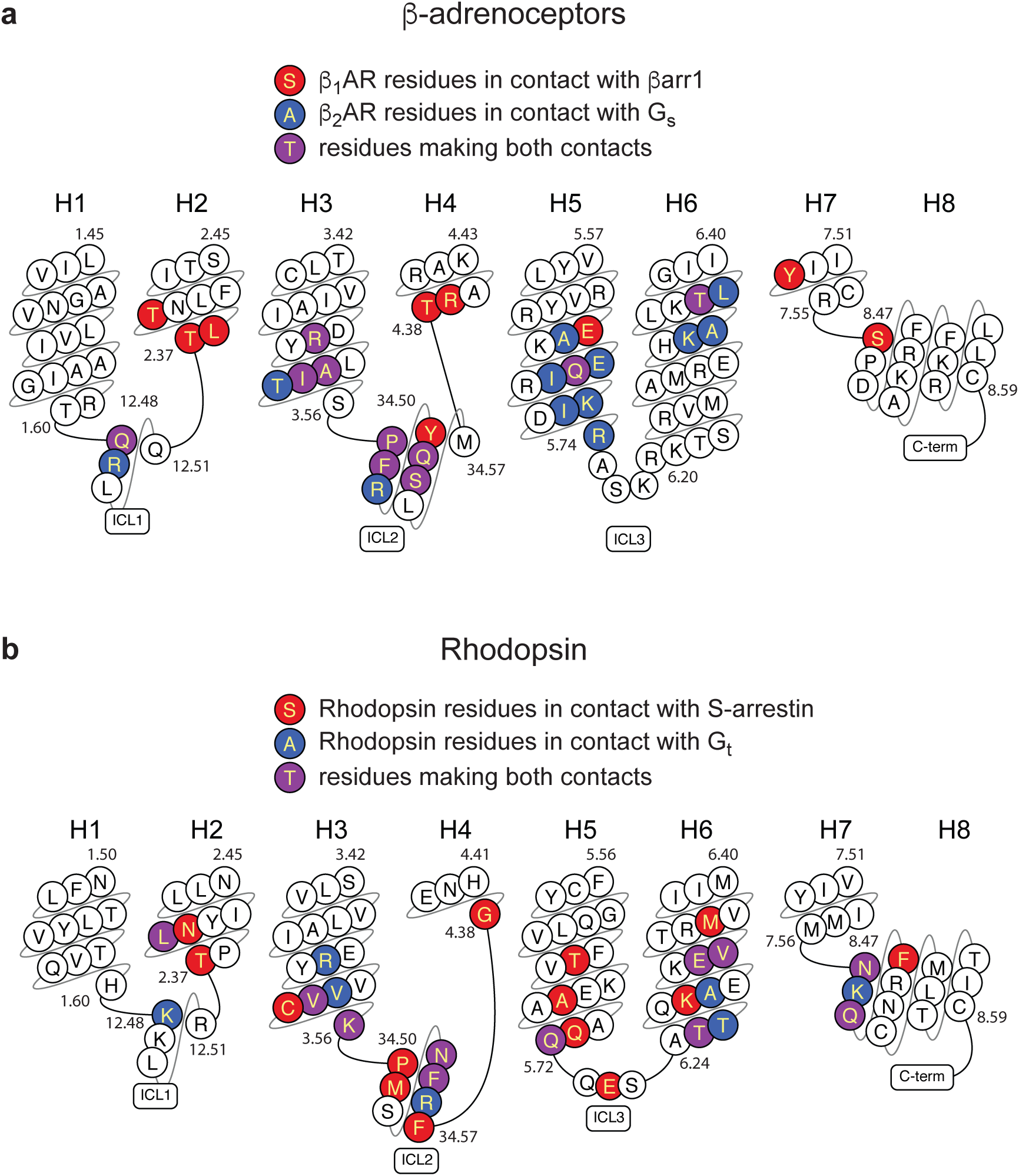
Comparison of the receptor-G protein and receptor-arrestin binding interface. Residues in GPCRs that make contact (within 3.9 Å) of arrestins or G proteins are highlighted. **a**, sequence of turkey β_1_AR is depicted. **b**, sequence of human rhodopsin is depicted. Plots were made using GPCRdb.

**Extended Data Fig. 8.**
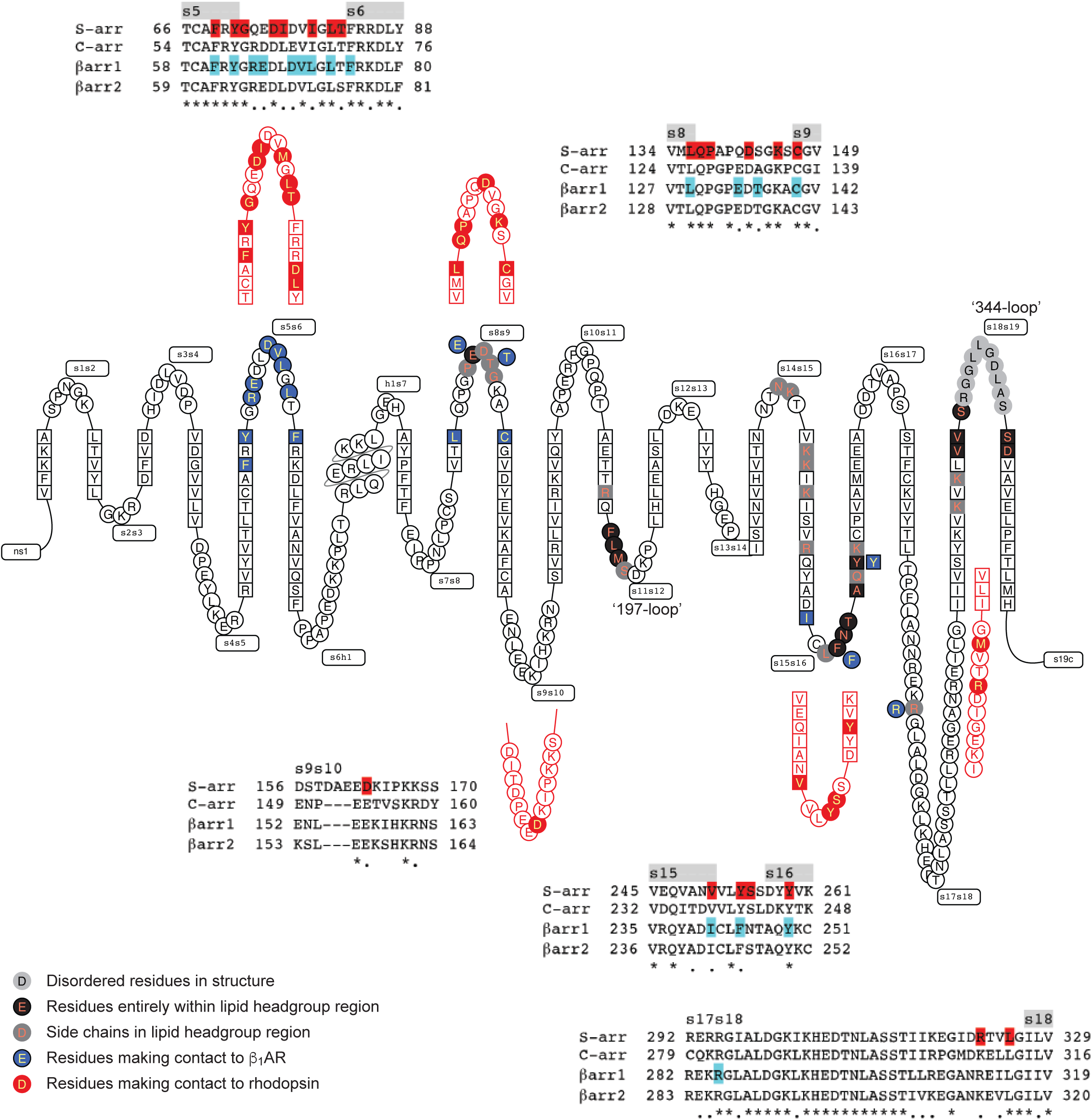
Comparison of the S-arrestin and βarr1 interfaces with GPCRs. A snake plot (GPCRdb) of human βarr1 depicts the secondary structure elements in the protein, with amino acid residues making contact with β_1_AR coloured blue. Equivalent regions in murine S-arrestin that make contact to rhodopsin are shown in red. Alignments of human arrestins show the variation of amino acid sequences within these specific regions, with residues making contact to the respective receptors highlighted.

**Extended Data Fig. 9.**
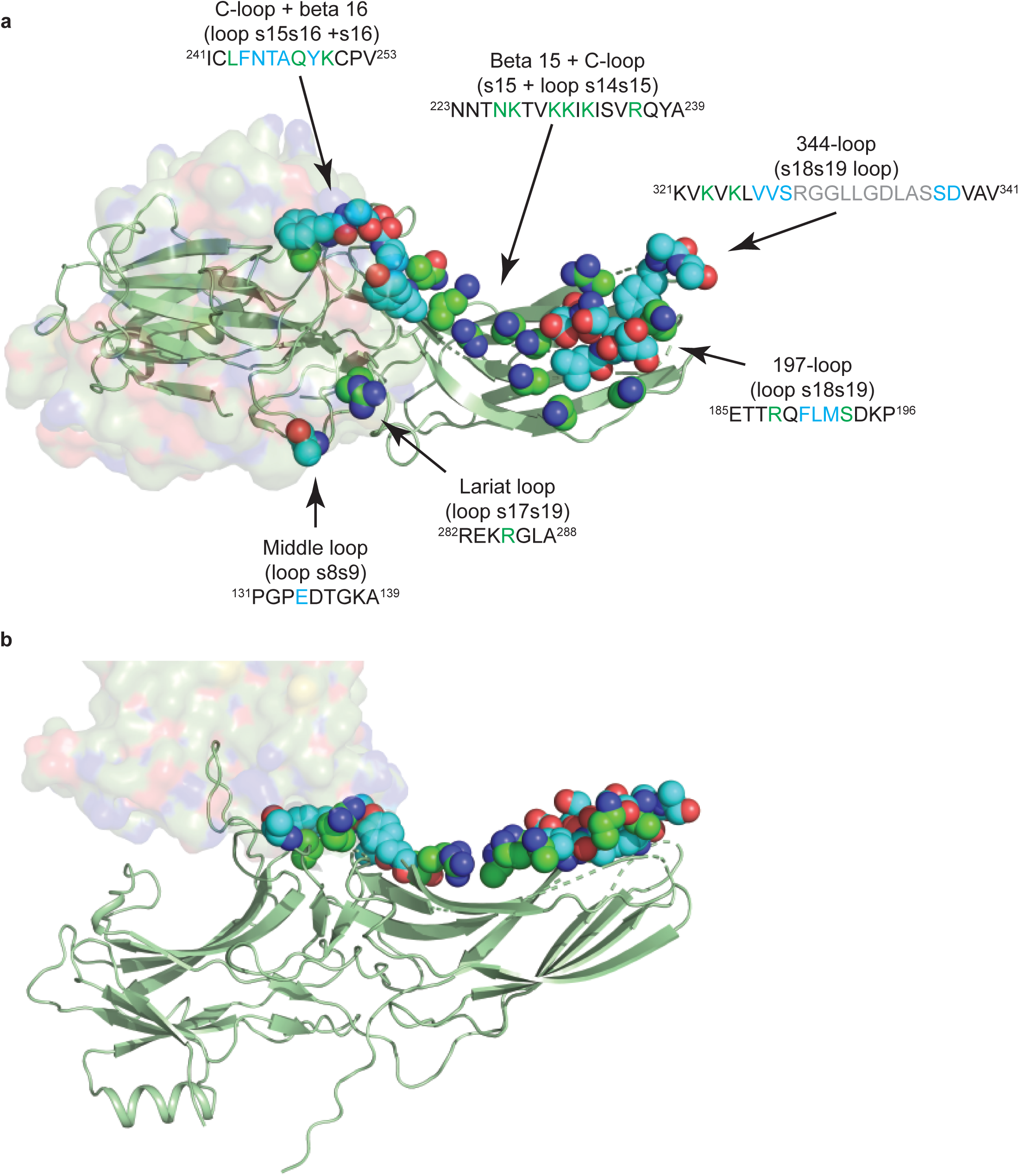
Lipid-interacting residues in βarr1. β_1_AR is depicted in surface representation and βarr1 as a cartoon (green) with atoms predicted to be within the head group region of the lipid bilayer shown as spheres: oxygen, red; nitrogen, blue; carbon, green or cyan. Residues coloured cyan are predicted to be entirely within the lipid head group region, whilst the carbons coloured green are the portions of these side chains that are potentially interacting with lipid head groups. **a**, view of the lipid interacting surface viewed through the receptor; **b**, view parallel to the membrane plane.

**Extended Data Fig. 10.**
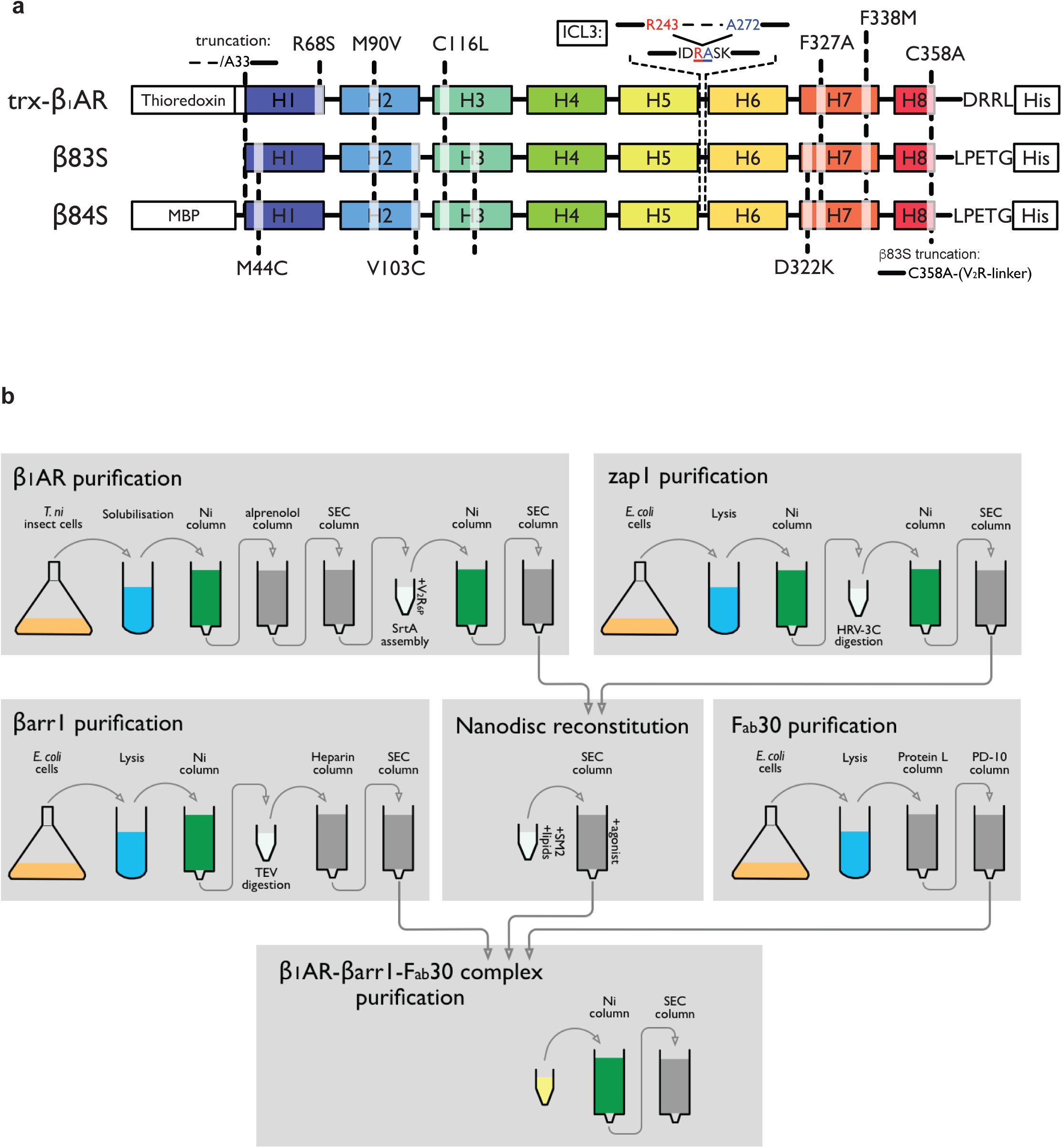
Description of constructs used for structural studies and the purification strategy. **a**, cartoon of the constructs used for X-ray crystallography (trx-β_1_AR), cryo-EM (β83S) and pharmacology (β84S) indicating the sites of truncations and point mutations. **b**, Purification scheme for the preparation of phosphorylated β_1_AR coupled to βarr1 for structure determination by cryo-EM.

**Extended Data Table 1.** Cryo-EM data collection and refinement statistics.

| | formoterol-bound $\beta_1$AR-β-arrestin-1-Fab30<br>(EMDB-10515)<br>(PDB 6TKO) | | |
| --- | --- | --- | --- |
| Session | LMB Krios2-1 | LMB Krios2-2 | DLS eBIC Krios1 |
| **Data collection and processing** |  |  |  |
| Magnification | 105,000x | 105,000x | 130,000x |
| Voltage (kV) | 300 | 300 | 300 |
| Electron exposure (e<sup>-</sup>/Å<sup>2</sup>) | 51 | 49 | 45 |
| Defocus range (μm) | -1.2 to -3.0 | -1.2 to -3.0 | -1.2 to -3.0 |
| Pixel size (Å) | 1.1 | 1.1 | 1.047 |
| Symmetry imposed | C1 | C1 | C1 |
| Initial particle images<sup>a</sup> (no.) | 2,257,195 | 1,959,236 | 448,633 |
| Contribution to final particle images (no.) | 175,204 | 183,140 | 45,647 |
| Final particle images (no.) |  | 403,991 |  |
| Map resolution (Å) |  | 3.3 |  |
| FSC threshold |  | 0.143 |  |
| Map resolution range<sup>b</sup> (Å) |  | ~3.2 to ~4.5 |  |
| **Refinement** |  |  |  |
| Initial model used (PDB code) |  | 6IBL, 4JQI |  |
| Model resolution<sup>c</sup> (Å) |  | 3.66 |  |
| FSC threshold |  | 0.5 |  |
| Map sharpening *B* factor (Å<sup>2</sup>) |  | -80 |  |
| Model composition |  |  |  |
| Non-hydrogen atoms |  | 8085 |  |
| Protein residues |  | 1052 |  |
| Ligands |  | 25 |  |
| *B* factors (Å<sup>2</sup>) |  |  |  |
| Protein |  | 80.5 |  |
| Ligand |  | 115 |  |
| R.m.s. deviations |  |  |  |
| Bond lengths (Å) |  | 0.001 |  |
| Bond angles (°) |  | 0.411 |  |
| Validation |  |  |  |
| Molprobity score |  | 1.06 |  |
| Clashscore |  | 2.75 |  |
| Poor rotamers (%) |  | 0.23 |  |
| EMRinger score |  | 2.06 |  |
| Ramachandran plot |  |  |  |
| Favored (%) |  | 99.02 |  |
| Allowed (%) |  | 0.98 |  |
| Disallowed (%) |  | 0 |  |

<sup>a</sup>After 3D classification.<sup>b</sup>Local resolution range.<sup>c</sup>Resolution at which FSC between map and model is 0.5.

**Extended Data Table 2.** X-ray data collection and refinement statistics.

| | formoterol-bound $\text{trx-}\beta_1\text{AR-Nb80}$<br>(PDB 6IBL) |
| --- | --- |
| **Data statistics** |  |
| Number of crystals | 1 |
| Space group | P 2<sub>1</sub> 2<sub>1</sub> 2<sub>1</sub> |
| Cell dimensions *a*, *b*, *c* (Å) | 116.6, 121.1, 129.8 |
| Resolution range (Å) | 44.28-2.7 (2.79-2.7)<sup>a</sup> |
| Unique reflections | 50611 (4416)<sup>a</sup> |
| Completeness before truncation (%) | 99.0 (95.3)<sup>a</sup> |
| Multiplicity | 4.6 (4.7)<sup>a</sup> |
| Mean I/σI | 6.6 (1.7)<sup>a</sup> |
| R<sub>merge</sub> | 0.151 (0.936)<sup>a</sup> |
| CC<sub>1/2</sub>=0.3 h, k, l axes & *overall* (Å) | 2.7, 3.62, 3.43, **2.92** |
| **Refinement statistics** |  |
| Resolution (Å) | 88.6-2.7 (2.78-2.7)<sup>a</sup> |
| Completeness, truncated data (%) | 62.71 (2.78) |
| No. of reflections | 30378 |
| R<sub>work</sub>/R<sub>free</sub> (%) | 0.242/0.276 (0.323/0.348)<sup>a</sup> |
| No. of atoms | 8271 |
| Protein | 7984 |
| Ligands & detergents | 258 |
| Water | 29 |
| B-factors (Å<sup>2</sup>) |  |
| Protein | 70.2 |
| Ligand & detergents | 57.9, 73.8 |
| Waters | 38.5 |
| R.M.S.D. |  |
| Bond lengths (Å) | 0.008 |
| Bond angles (°) | 1.17 |

<sup>a</sup>Outer resolution shell.

